# Bi-cross-validation: a data-driven method to evaluate dynamic functional connectivity models in fMRI

**DOI:** 10.64898/2026.04.02.716067

**Authors:** Yiming Wei, Stephen M. Smith, Chetan Gohil, Rukuang Huang, Ben Griffin, SungJun Cho, Stanislaw Adaszewski, Stefan Fraessle, Mark W. Woolrich, Seyedeh-Rezvan Farahibozorg

**Affiliations:** Centre for Integrative Neuroimaging, FMRIB, Nuffield Department of Clinical Neurosciences, University of Oxford, Oxford OX3 9DU, United Kingdom; Oxford Centre for Human Brain Activity (OHBA), Centre for Integrative Neuroimaging, Department of Psychiatry, University of Oxford, Oxford, OX3 7JX, United Kingdom; Computational Sciences Center of Excellence, F. Hoffmann-La Roche Ltd., CH-4070 Basel, Switzerland

**Keywords:** dynamic functional connectivity, brain connectivity, resting-state fMRI, model selection, cross-validation

## Abstract

Dynamic functional connectivity (dFC) models have become increasingly popular over the past decade for characterising time-varying interactions between brain regions. However, assessing and comparing dFC models remains challenging. Here, we introduce bi-cross-validation as a general framework for evaluating dFC models and selecting key hyperparameters, such as the number of states. By jointly partitioning the data across subjects and brain regions, bi-cross-validation enables out-of-sample evaluation without re-estimating latent states on the same data used for testing, thereby avoiding circularity. Using simulated data with known ground-truth dynamics, we show that bi-cross-validation favours models that accurately capture the underlying state structure. Applying the framework to real resting-state fMRI data, we demonstrate that bi-cross-validation naturally balances goodness-of-fit against model complexity, with performance improving and then declining as model complexity increases. Finally, we use bi-cross-validation to directly compare static and dynamic FC models, showing that dynamic models underperform static models at low spatial dimensionality, but outperform static models at sufficiently high dimensionality. Together, these results establish bi-cross-validation as a principled tool for dFC model selection, evaluation, and comparison.

## 1 Introduction

Functional connectivity (FC) quantifies interactions between brain regions, commonly defined as the correlation between their neurophysiological time series (Friston, 1994), and is often studied using functional magnetic resonance imaging (fMRI) and electrophysiological modalities such as magnetoencephalography (MEG) and electroencephalography (EEG). Conventional FC is estimated using a static approach that calculates a single correlation value between each pair of brain regions across the entire scanning session. In recent years, there has been growing interest in dynamic functional connectivity (dFC), which refers to temporal fluctuations in FC on the order of seconds to minutes (Calhoun et al., 2014; Chang & Glover, 2010; Hutchison et al., 2013; Lurie et al., 2020). Since brain activity is inherently dynamic, time-varying network descriptions may capture information that static approaches overlook, potentially revealing fundamental principles of brain organisation. Dynamic functional connectivity has been shown to provide potential biomarkers of ongoing cognition (such as consciousness and daydreaming) (Barttfeld et al., 2015; Kucyi & Davis, 2014), cognitive and behavioural traits (Liégeois et al., 2019; Vidaurre et al., 2017), and psychiatric conditions (Kaiser et al., 2016; Sakoglu et al., 2010).

To capture these dynamics, various models have been developed. The sliding window correlation (SWC) method divides a scanning session into fixed-length windows and computes the average correlation within each window (Sakoglu et al., 2010). Functional connectivity matrices from different windows are then clustered using algorithms such as k-means to identify recurring, transient connectivity patterns (Allen et al., 2014). A key challenge in SWC is that the window length must be long enough to ensure reliable estimation of FC (Hutchison et al., 2013), yet short enough to detect rapid transitions. To address this limitation, the Hidden Markov Model (HMM) has been applied to characterise dFC through the automated detection of transient brain states (Vidaurre et al., 2017). Conceptually, it models the data as arising from a sequence of latent states with distinct connectivity patterns, estimating both the states themselves and their probabilistic transitions. Recent models, such as Dynamic Network Modes (DyNeMo) (Gohil et al., 2022) and Multi-dynamic Adversarial Generator Encoder (MAGE) (Pervaiz et al., 2022), extend beyond the HMM by modelling mixtures of modes at each time point^1^ and incorporating long-range temporal dependencies using deep learning techniques. While many dFC methods characterise whole-brain temporal dynamics in the time domain, other approaches focus on the frequency domain (Chang & Glover, 2010; Yaesoubi et al., 2015), or analyse subsets of time points, such as co-activation patterns (CAPs) (Liu & Duyn, 2013; Liu et al., 2018). Although most studies have focused on restingstate fMRI, these methods are also applicable to task fMRI (Gonzalez-Castillo & Bandettini, 2018) and electrophysiological modalities such as MEG (Baker et al., 2014; Gohil et al., 2022).

Evaluating and optimising dFC models remains challenging (Lurie et al., 2020). At their core, these models are a form of unsupervised learning: they explain fMRI time series by identifying latent “states” or “clusters” with distinct connectivity patterns, but the true states underlying the data are unknown. In such settings, an effective model should balance goodness-of-fit with model complexity. Goodness-of-fit is relatively straightforward to quantify, for example by evaluating how well the model explains observed data (e.g., log-likelihood in HMM and DyNeMo (Gohil et al., 2022; Vidaurre et al., 2017)), or how closely sliding-window FC matrices match cluster centroids (cluster validity index in SWC with k-means (Allen et al., 2014)). Model complexity, however, is harder to assess: existing penalty terms within each model framework often perform poorly, and there is no principled way to compare complexity across different models. Cross-validation may seem a natural solution. However, in unsupervised learning it becomes problematic because evaluation on a validation set requires inferring hidden states from that same set, making the evaluation inherently circular.

To address these limitations, we propose bi-cross-validation (Fu & Perry, 2017; Owen & Perry, 2009) as a principled technique to evaluate dFC models and optimise their hyperparameters. Originally developed for unsupervised learning tasks such as matrix decomposition (Owen & Perry, 2009) and k-means clustering (Fu & Perry, 2017), bi-cross-validation naturally generalises to dFC model evaluation by simultaneously splitting data along the “subject” and “brain region” dimensions. Following a four-step validation procedure (described in Methods), latent states or clusters are inferred from one subset of regions in the validation set, while the scoring function is computed using the complementary subset, thus circumventing the circularity issue. Importantly, this approach enables principled comparisons across different dFC models, provided they offer a probabilistic description of whole-brain fMRI data.

The bi-cross-validation framework integrates and extends previous evaluation efforts in three ways, each demonstrated in the main results section:

### 1. Assessing model sensitivity with simulated data

Researchers simulate fMRI data with known transitions to assess how effectively models detect ground-truth changes in functional connectivity (Hindriks et al., 2016; Lindquist et al., 2014; Shakil et al., 2016; Thompson et al., 2018). However, the generalisability of simulation-based evaluations to real fMRI data remains uncertain. In our simulations (Section 3.1), we show that bi-cross-validation favours models aligned with the true state structure.

### 2. Balancing goodness-of-fit and model complexity on real data

Previous studies have quantified model complexity indirectly, often through reproducibility or test–retest reliability, metrics that typically decrease monotonically as model complexity increases (Abrol et al., 2017; Choe et al., 2017; Zhang et al., 2018). Yet, explicitly balancing complexity against goodness-of-fit remains challenging, as these aspects are typically evaluated separately. Using real resting-state fMRI data (Section 3.2), we demonstrate that bi-cross-validation naturally achieves this balance: model performance initially improves but eventually declines as complexity (number of states) increases. We further illustrate this advantage by comparing bi-cross-validation directly against previously proposed metrics of model fit and reproducibility.

### 3. Comparing static and dynamic FC models

Determining whether observed fluctuations in FC represent genuine changes in network organisation or merely sampling variability remains an open question (Laumann et al., 2017), with existing null-model-based approaches producing ambiguous or inconclusive results (Liegeois et al., 2017, 2021). Bi-cross-validation reframes this issue as a model comparison problem. Specifically, by setting the number of states to one, it evaluates static FC models against multi-state dynamic models, directly testing for the presence of meaningful temporal dynamics (Section 3.4).

## 2 Methods

In this section, we describe the dFC models evaluated in this study (Section 2.1; Figure 1), outlining their shared methodological framework and fitting procedures. We then introduce our bi-cross-validation approach (Section 2.2; Figure 2) and compare it with conventional evaluation metrics (Section 2.3). Finally, we provide details of simulation paradigms and real fMRI datasets used for validation (Section 2.4).

**Figure 1.**
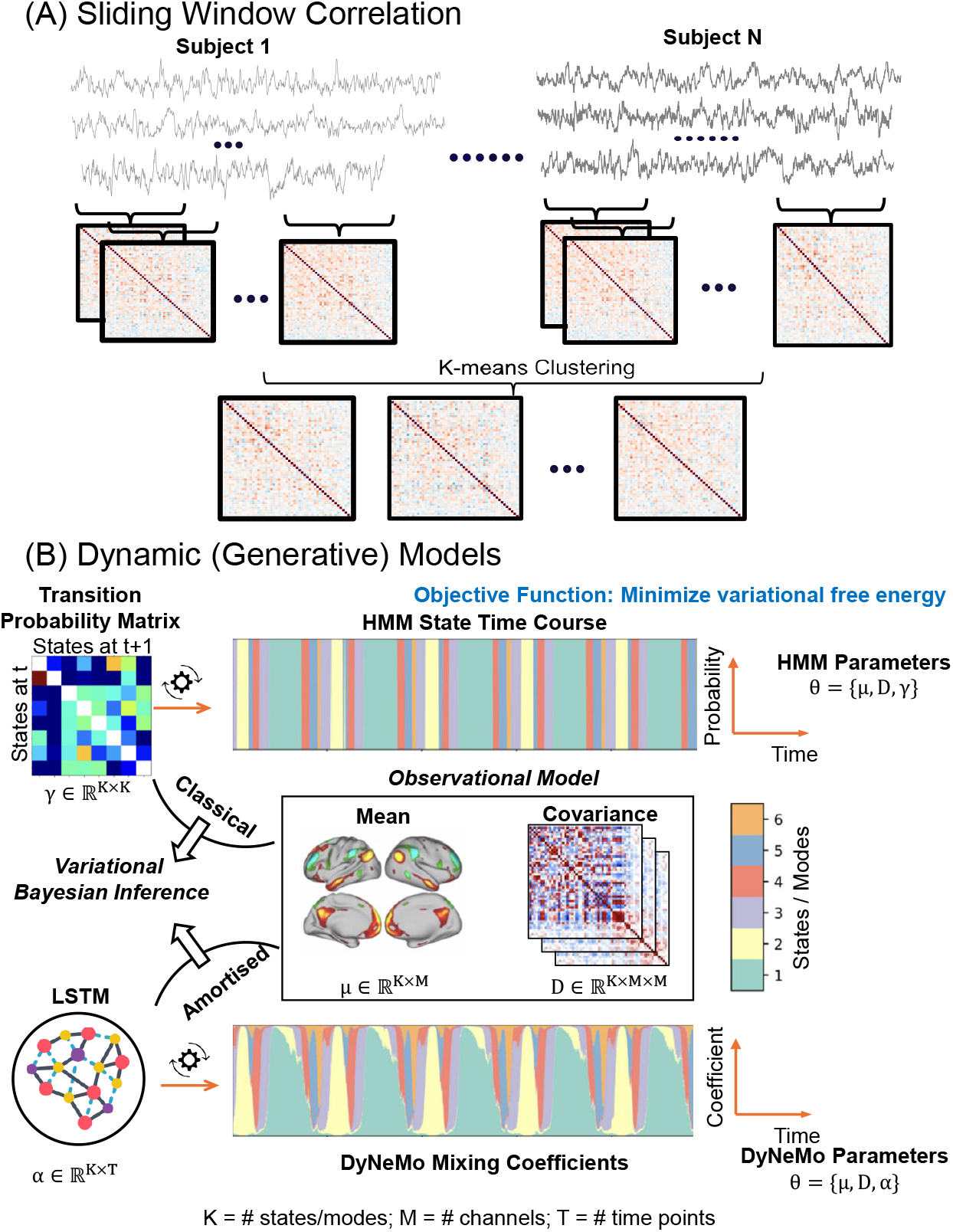
Dynamic functional connectivity models. (A) Sliding window correlation (SWC). SWC divides the scanning session into overlapping or non-overlapping windows and computes functional connectivity within each window. A clustering algorithm, such as k-means, is commonly applied to identify recurring patterns of functional connectivity. (B) Hidden Markov Model (HMM) and Dynamic Network Modes (DyNeMo). The HMM assumes that brain activity transitions between a finite set of discrete states, each characterised by a distinct mean activation pattern and covariance structure. These transitions are governed by a transition probability matrix. Model parameters are estimated using variational Bayesian inference. DyNeMo extends the HMM in two key ways: (1) it models brain activity as a continuous mixture of “modes” rather than discrete states, with each mode defined by its own mean activation and covariance matrix; and (2) it incorporates long-range temporal dependencies using a Long Short-Term Memory (LSTM) network, which models the dynamic mixing of modes over time. DyNeMo parameters are inferred using amortised variational Bayesian inference.

**Figure 2.**
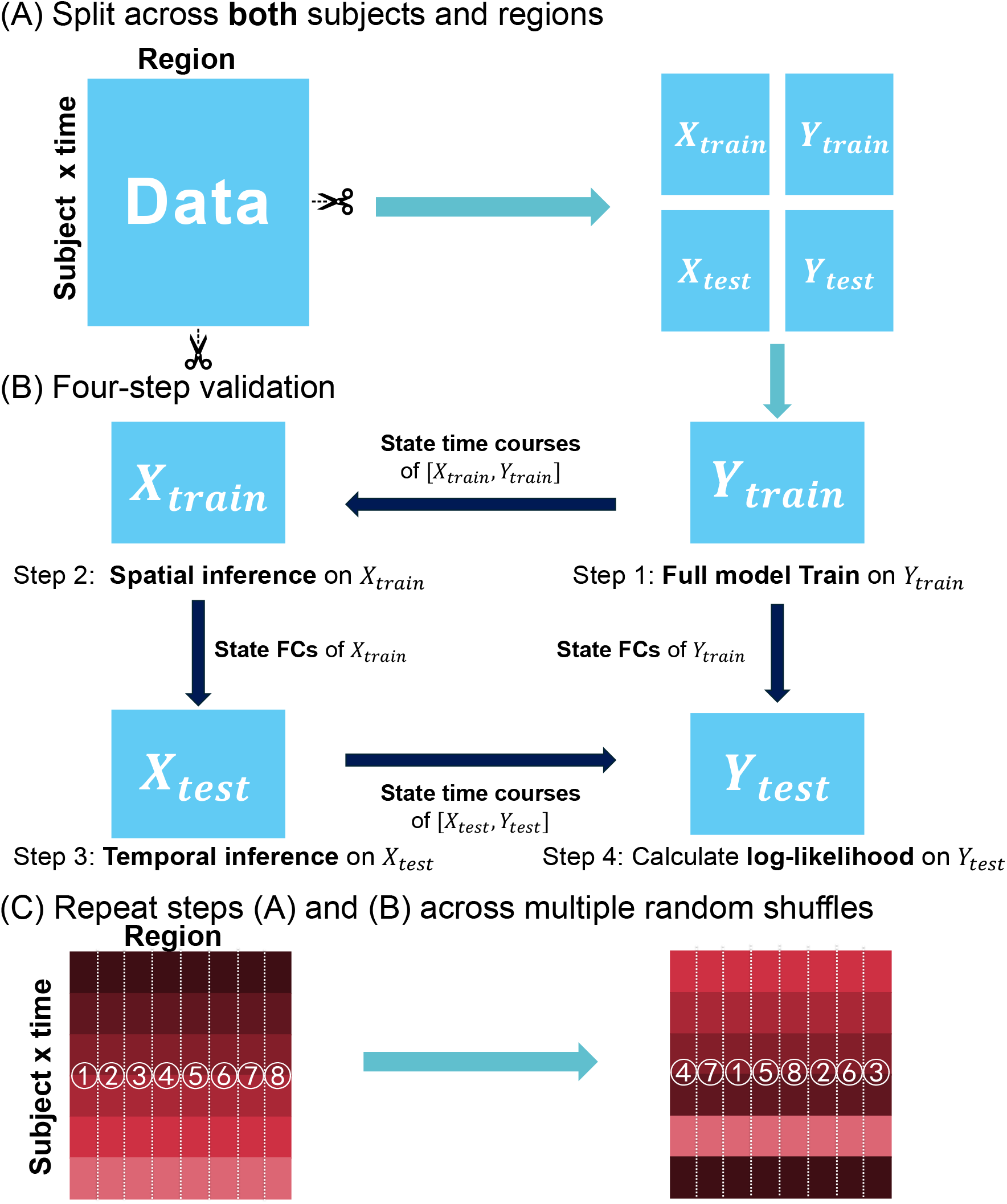
Bi-cross-validation. (A) After concatenation across subjects, the fMRI data are partitioned along both rows (subjects × time) and columns (regions), yielding four data blocks: [*X*_train_, *Y*_train_; *X*_test_, *Y*_test_]. (B) The resulting data blocks undergo a four-step evaluation procedure. Step 1: A full model is trained on *Y*_train_, estimating the state time courses, observation model parameters, and temporal dynamics. Step 2: The state time courses estimated from *Y*_train_ are reused to infer observation model parameters on *X*_train_ (spatial inference). Step 3: The inferred observation model parameters from *X*_train_ are fixed to infer state time courses on *X*_test_ (temporal inference). Step 4: The observation model parameters estimated in Step 1 and the state time courses from Step 3 are combined to compute the log-likelihood on *Y*_test_. (C) The procedure is repeated across multiple random shuffles of the data, and performance is aggregated across repetitions.

### 2.1 Models evaluated

#### 2.1.1 Sliding Window Correlation (SWC)

The sliding window correlation (SWC) method estimates dFC by applying a sliding window to fMRI time series and computing an FC matrix within each window (Figure 1A) (Sakoglu et al., 2010). These matrices are then clustered to identify recurring connectivity patterns, or “states” (Allen et al., 2014). While several methodological choices, such as window type, window length, and window offset, can influence performance (Lurie et al., 2020), we focus on optimising the number of clusters/states *N*_states_, while keeping other parameters fixed. The key implementation steps are as follows:

- **Windowing**: Human Connectome Project (Smith et al., 2013) and simulated data (1200 time points): 100-TR rectangular window, 100-TR offset (12 windows per session). UK Biobank data (490 time points) (Miller et al., 2016): 70-TR rectangular window, 70-TR window offset (7 windows per session). Here, TR refers to the repetition time, i.e. the temporal resolution of fMRI, and the offset specifies the step size between successive windows.
- **Clustering**: The lower-triangular elements of each windowed FC matrix were vectorised and clustered using k-means to derive *N*_states_ centroids. Although the number of windows per subject is modest, clustering was performed across all subjects, yielding a large number of windowed FC samples overall and ensuring stable estimation of state centroids.

We analysed state covariances rather than correlations. Because z-scoring is applied to each regional time series at the full-session level, variance can still vary across windows; covariances thus retain information on both fluctuation magnitude and coupling between regions. This implementation prioritises evaluating the effect of the number of states, rather than exhaustively optimising all parameters.

#### 2.1.2 Static Functional Connectivity (sFC)

A special case of SWC occurs when the window spans the entire session, reducing the method to static functional connectivity (sFC). In this case, a single covariance matrix was estimated for each subject. To maintain comparability with the dynamic models, these subject-level covariance matrices were clustered across subjects using k-means to capture inter-subject variability. This clustered sFC model serves as a baseline and allows us to assess whether improvements in dFC models arise from genuine temporal dynamics, subject variability, or both.

#### 2.1.3 Hidden Markov Model (HMM)

The Hidden Markov Model (HMM) provides an instantaneous estimation of FC by assigning a distinct brain state to each time point, in contrast to the windowed approach used in SWC (Figure 1B) (Baker et al., 2014; Vidaurre et al., 2017). This time-point level mapping enables the detection of fast-evolving connectivity dynamics that may be missed by window-based methods. The model imposes temporal regularisation by assuming Markovian dynamics, where transitions between a finite set of states are governed by a transition probability matrix. The probability of transitioning to a new state depends only on the immediately preceding state. Each state is characterised by a multivariate Gaussian distribution. A key hyperparameter in the HMM is the number of states. Too few states may fail to capture the complexity of the data, while too many can lead to overfitting. Ideally, the optimal number of states balances goodness-of-fit and model complexity. Determining this number has been a long-standing challenge in neuroimaging, and a common approach is to fit the HMM across multiple candidate numbers of states (Griffin et al., 2024; Vidaurre et al., 2017).

#### 2.1.4 Dynamic Network Modes (DyNeMo)

Dynamic Network Modes (DyNeMo), proposed by Gohil et al. (2022), generalises the HMM by replacing discrete, mutually exclusive states with a continuous mixture of “modes” (Figure 1B). Each mode is characterised by a multivariate Gaussian distribution, and functional connectivity at each time point is modelled as a weighted linear combination of these modes. The time-varying normalised weights define the dynamic contribution of each mode, offering a more flexible representation of brain dynamics than the discrete-state assumption in the HMM. To model the temporal evolution of these weights, DyNeMo employs a Long Short-Term Memory network (LSTM) (Hochreiter & Schmidhuber, 1997), which captures long-range temporal dependencies. Like the HMM, DyNeMo requires selecting the number of modes, leading to a similar hyperparameter tuning challenge.

#### 2.1.5 Model summary

Our evaluation framework is grounded in the idea that dFC models should provide a low-dimensional representation of fMRI data in the time domain. Within each session, the preprocessed fMRI dataset typically consists of a large matrix with hundreds of time points and tens of regions. dFC models aim to re-express this matrix using a small number of states or modes, represented through state time courses (or linear coefficients) and associated state/mode statistics. In the HMM, latent states are mutually exclusive, such that exactly one state is active at each time point, with transitions governed by a transition probability matrix. In DyNeMo, the time courses are continuous mixing coefficients governed by a recurrent neural network, constrained to sum to one at each time point. In SWC, state time courses correspond to the window labels derived from k-means clustering. Notably, when the number of states/modes is set to one, all models reduce to a group-level static FC model. A summary of model parameters is provided in Table 1.

**Table 1:** Summary of model parameters. The state/mode time courses represent the brain state (or additive combination of modes) at each time point. The observation model parameters incorporate covariance statistics for each state/mode, while the temporal dynamics parameters govern the evolution of state/mode time courses over time. In HMM and DyNeMo inference (see Section 2.1.6), `osl-dynamics` estimates an approximate distribution for the state/mode time courses and provides a point estimate for both the observation model and temporal dynamics parameters. In contrast, the SWC follows a heuristic fitting procedure, yielding only point estimates of all these parameters.

| Model | State time courses | Observation model | Temporal dynamics |
| --- | --- | --- | --- |
| HMM | binary | State covariances | Transition probability matrix |
| DyNeMo | non-binary | Mode covariances | LSTM |
| SWC | binary (window label) | Centroid covariances | – |

#### 2.1.6 Model inferences

The inference processes for HMM and DyNeMo aim to estimate the posterior distribution of unobserved variables given the observed data. In most cases including this study, obtaining the full posterior distribution over these variables is computationally infeasible. To address this, we use the variational Bayesian method, which approximates the true posterior with a more tractable surrogate distribution (Beal, 2003). To simplify the problem further, we assume that point estimates are sufficient for all model parameters except for the state time courses or mode linear coefficients. Mathematically, the variational Bayesian method optimises a loss function known as variational free energy using stochastic gradient descent. A detailed mathematical description of the inference procedure is provided by Gohil et al. (2023) and Gohil et al. (2022). All generative model inference procedures were implemented using the `osl-dynamics` software package (Gohil et al., 2023). The specific inference settings used in this study are provided in Supplementary Tables S1 and S2.

In contrast, SWC does not require advanced inference techniques. It estimates sample covariances within each window and subsequently clusters these matrices using the k-means algorithm. The resulting cluster labels define the state time courses (i.e., the assignment of windows to states), and the cluster centroids represent the observation model parameters.

Regardless of the specific model or inference technique, each method involves estimating three main components: the state time courses, observation model parameters, and temporal dynamics parameters (Table 1). When all parameters are inferred jointly, we refer to this as full model training. Crucially, our evaluation framework builds on two additional inference modes in which subsets of parameters are held fixed:

1. **Spatial Inference**: The state time courses and temporal dynamics parameters are fixed, and only the observation model parameters are estimated. This approach focuses on refining the spatial patterns (e.g., covariance structure) of each state or mode.
2. **Temporal Inference**: The observation model parameters are fixed while the state time courses and temporal dynamics parameters are estimated. This approach is concerned with capturing the temporal evolution of the states or modes.

### 2.2 Bi-cross-validation

#### 2.2.1 Intuition

Cross-validation, widely used in supervised learning, assesses performance on a held-out validation set (Hastie et al., 2009). However, its direct application to unsupervised learning problems, such as singular value decomposition (SVD), k-means clustering, and dFC models, is not straightforward. A standard approach might involve training a full HMM on the training set, then inferring the state time courses on the validation set and computing goodness-of-fit metrics such as free energy or log-likelihood. However, this procedure inadvertently incorporates information from the validation set before the evaluation step, violating a core principle of cross-validation. As a result, overfit models can create spurious states that capture noise in the data. During validation, these states are reassigned to match noisy fluctuations in the held-out set, which artificially inflates apparent performance. Because standard cross-validation does not impose an explicit complexity penalty in this setting, such models are not appropriately penalised for overfitting.

To address this issue, Fu and Perry (2017) and Owen and Perry (2009) introduced bi-cross-validation, a technique designed for unsupervised model evaluation. The key idea is to reshuffle and split **both** the samples and variables of the dataset (Figure 2A). In this framework, model performance is assessed by inferring latent structure (e.g., labels or time courses) using a subset of the variables within the validation set and then computing prediction error on the remaining variables. By separating inference and evaluation across different subsets of variables, bi-cross-validation prevents information from leaking between the two steps and, crucially, penalises overfitting: spurious states that fit only idiosyncratic noise in one subset of variables will fail to generalise to the held-out subset. This provides an implicit check on model complexity that standard cross-validation lacks. Originally developed for SVD (Owen & Perry, 2009) and k-means clustering (Fu & Perry, 2017), here we adapt bi-cross-validation to evaluate dFC models.

#### 2.2.2 Details

We first concatenate all subjects’ data along the time axis, yielding a data matrix of shape (*N*_subjects_ × *N*_timepoints_, *N*_regions_) (Figure 2A). Rows correspond to time points from individual subjects, and columns correspond to brain regions. Bi-cross-validation partitions this matrix along both dimensions to produce four data blocks: [*X*_train_, *Y*_train_; *X*_test_, *Y*_test_] (Figure 2A). Because fMRI time series exhibit strong temporal dependence, we do not split within a subject’s time series. Instead, all time points from a given subject are assigned entirely to either the training or the test set. This ensures independence between training and test samples and prevents data leakage (Rosenblatt et al., 2024).

Bi-cross-validation then proceeds through a four-step evaluation procedure (Figure 2B); a full mathematical description is provided in Appendix A.

1. Train a full model on *Y*_train_ to estimate state time courses, observation model parameters and temporal dynamics.
2. Use the state time courses from *Y*_train_ to infer observation model parameters for *X*_train_ (spatial inference).
3. Use the observation model parameters estimated from *X*_train_ to infer the state time courses on *X*_test_ (temporal inference).
4. Combine the observation model parameters from *Y*_train_ and the state time courses from *X*_test_ to compute model performance on *Y*_test_.

Final model evaluation is always performed on *Y*_test_.

Because the bi-cross-validation score depends on the specific partition of rows and columns, relying on a single split would yield an estimate sensitive to arbitrary partitioning. Therefore, prior to partitioning, the rows and columns are randomly permuted, and the full procedure is repeated across multiple shuffles (Figure 2C).

#### 2.2.3 Remarks

Several implementation details are worth noting. First, we use log-likelihood as the final scoring function because both HMM and DyNeMo explicitly assume multivariate Gaussian observation models (Gohil et al., 2022; Vidaurre et al., 2017), and SWC is commonly estimated under the same assumption (Hindriks et al., 2016). For simplicity, we assume zero-mean Gaussian observations that are independent and identically distributed within each latent state or mode, consistent with prior fMRI applications of HMM (Vidaurre et al., 2017). Temporal dependencies are captured by the latent dynamics through Markov transitions in the HMM and recurrent dynamics in DyNeMo. The framework can readily be extended to richer observation models, including non-zero mean activation, autoregressive observation processes (Baker et al., 2014), haemodynamic response modelling (Lindquist et al., 2009), and subject-specific embeddings (Huang et al., 2026).

Second, bi-cross-validation assumes that brain states (or modes) involve multiple regions and that time series from different brain areas are not perfectly correlated. If states are specific to individual regions, the method would fail; if brain signals were perfectly correlated, it would reduce to standard cross-validation because any noise overfitting in *Y* would simply be propagated through *X* and back to *Y*, leaving the overfitting problem unresolved. A linear transformation, as suggested in Fu and Perry (2017), could mitigate such dependencies. However, given the typically low correlation among brain regions and the widespread spatial distribution of dFC states (at least, when given a sufficiently fine-grained parcellation), we consider this step unnecessary in our setting.

Third, although k-fold cross-validation is common in supervised learning, we used a 1:1 split between training and test samples, and likewise a 1:1 split of regions into *X* and *Y*. This balanced partition improves the stability of parameter estimation in bi-cross-validation. To obtain a robust performance estimate, we generated 100 random splits using the `ShuffleSplit` class from `scipy` and applied the four-step procedure to each split for all models and hyperparameter settings.

Fourth, when *N*_states_ = 1, all models reduce to a group-level “static” FC model. In this case, spatial and temporal inference (steps 2 and 3) are unnecessary (Figure 2B). We estimate a single covariance matrix from *Y*_train_ and compute log-likelihood on *Y*_test_. To assess whether dFC improves model fit, we subtract the log-likelihood of the static model (with *N*_states_ = 1) from that of each dynamic model (*N*_states_ *>* 1) under the same split. This centring procedure helps visualise performance improvements due to dynamics.

Fifth, although model performance is generally expected to exhibit a peak as a function of model order, in practice we often observe a plateau near the optimum. This can occur because some models effectively down-weight redundant states or modes, so additional capacity does not substantially increase effective complexity. Moreover, neighbouring model orders may yield statistically indistinguishable fits, resulting in a flat region around the optimum. While users may adopt different criteria for model selection, we follow a two-step procedure. We first identify the model (i.e., number of states) with the highest median of bi-cross-validated log-likelihood across all realisations. We then perform a backward search, reducing the number of states or modes until performance becomes significantly worse than that of the best model. Significance is assessed using a one-sided paired t-test with a threshold of *p* = 0.001. In our experience, the selected model order is robust to moderate changes in this threshold.

Sixth, an alternative masking strategy used in earlier work is to randomly mask scattered entries of the data matrix and evaluate reconstruction error on those held-out elements (Wold, 1978). We do not adopt this strategy because it would disrupt the spatial and, critically, temporal dependencies present in fMRI data, and most dFC models are not designed to be trained with arbitrarily missing observations.

### 2.3 Conventional evaluation metrics

In addition to bi-cross-validation, we implemented several conventional evaluation metrics commonly used in the literature. While these metrics are not as complex as bi-cross-validation, they provide valuable insights into model behaviour and help validate the effectiveness of our proposed framework.

#### 2.3.1 Split-half reproducibility

Split-half reproducibility assesses whether the recurring connectivity patterns identified by a model are consistently detected across different subsets of the dataset. We followed the approach in previous studies (Gohil et al., 2022; Pervaiz et al., 2022), in which subjects were randomly divided into two equal halves, and the model was trained separately on each subset using the same hyperparameters. States (or modes) obtained from both splits were then paired using the following procedure:

1. After model training, we computed the pairwise Riemannian distance (Lim et al., 2019) between the state/mode covariance matrices from the two splits.
2. The Hungarian algorithm (Kuhn, 1955) was used to optimally match states/modes between splits.
3. The mean Riemannian distance between the matched states (or modes) served as the reproducibility metric.

To ensure robustness, this procedure was repeated five times with different random subject splits. Notably, bi-cross-validation implicitly penalises poor reproducibility: if the states or modes are inconsistent between *Y*_train_ and *Y*_test_ (or *X*_train_ and *X*_test_), the model will struggle to recover time courses in step 3 and generalise state covariances in step 4 (Figure 2B), ultimately resulting in reduced bi-cross-validation performance.

Reproducibility alone does not assess how well the model describes the data. A model may yield highly consistent states or modes across splits while providing a suboptimal fit. Therefore, it serves as a complementary metric rather than a comprehensive criterion for model selection.

#### 2.3.2 Standard cross-validation

As outlined in Section 2.2, standard cross-validation is problematic for evaluating dFC models. Nevertheless, we implemented a standard 5-fold cross-validation procedure to illustrate its shortcomings. Specifically, the model was trained on the training set, and state time courses (or mode linear coefficients) were inferred directly from the validation data. Model fit on the validation set was then quantified as variational free energy for HMM and DyNeMo, which reflects an approximate lower bound on the marginal likelihood, and as log-likelihood for SWC, quantifying how well the centroid covariances explained the windowed validation data.

#### 2.3.3 Fractional occupancy

Fractional occupancy is a post-hoc metric that quantifies the proportion of time spent in each state over the duration of the data. We computed this measure to assess whether the inferred states reflect genuine temporal dynamics shared across subjects, rather than being driven by a small subset of individuals or by modes with negligible contribution. Since the states or modes in all these models are defined at the group level, this diagnostic helps ensure that the model does not rely on states that are infrequently visited by most subjects. For HMM, we computed fractional occupancy for each subject and plotted the resulting distributions across states. Defining an analogous measure for DyNeMo is less straightforward, as modes can be non-exclusive and overlap in time. To characterise mode prominence in this setting, we used the standard deviation of the posterior mode time courses, normalised by corresponding mode covariances, as a measure of each mode’s contribution to time-varying functional connectivity.

### 2.4 Data Acquisition and Pre-processing

#### 2.4.1 Real data

We used resting-state fMRI data from the Human Connectome Project (HCP) S1200 release as our primary analysis dataset. Detailed information on data acquisition and pre-processing is available in Glasser et al. (2013) and Van Essen et al. (2013). The dataset comprised 1003 subjects, each of whom completed four 15-minute resting-state fMRI sessions with TR = 0.72 s. Additional pre-processing included ICA-FIX denoising (single-subject ICA followed by the FIX classifier) and between-subject spatial alignment using MSMAll, which utilises both structural and functional features (Robinson et al., 2014). Group-level spatial Independent Component Analysis (ICA) was subsequently applied following the method described in Smith et al. (2014), yielding spatial maps and time series for various numbers of independent components (*N*_regions_). We evaluated results with 15, 25, 50, and 100 components. All time series were z-scored for each component, session, and subject. Since low-frequency components can drive spurious correlations, particularly in SWC (Leonardi & Van De Ville, 2015), in some of the analyses, we applied a 5th-order Butterworth high-pass filter with a cut-off frequency of 0.05 Hz. When relevant, this is specified in the Results section.

We also analysed resting-state fMRI data from UK Biobank (UKB) (Miller et al., 2016) to evaluate the generalisability of our findings. Details of the automatic pre-processing pipeline are provided in Alfaro-Almagro et al. (2018). We randomly selected 5000 subjects from a pool of 70, 000 baseline scans. Each subject underwent a 6-minute resting-state scan with TR = 0.735 s. A key difference in the UKB pipeline is the application of conservative high-pass filtering prior to ICA-FIX denoising, using Gaussian-weighted least-squares straight-line fitting with *σ* = 50.0 s, to remove slow signal drift. To extract independent component time courses for each subject, the CIFTI-based rfMRI data were processed via dual regression using HCP group-level CIFTI spatial maps with 25 and 50 components. The resulting time series were then z-scored, and, as with the HCP data, were high-pass filtered at 0.05 Hz in some of the analyses. When relevant, this is specified in the Results section.

#### 2.4.2 Simulated data

We generated synthetic datasets to evaluate whether bi-cross-validation could accurately recover the underlying model and hyperparameter settings. Data were generated from the generative models underlying HMM and DyNeMo, each configured with six states or modes, 50 regions, and 1200 time points per subject, across 500 subjects. We did not generate simulations using sliding window correlation (SWC), as it does not specify an explicit generative model. All simulations were implemented using the `osl-dynamics` toolbox (Gohil et al., 2023).

To assess performance under realistic conditions, we conducted a set of simulations based on models trained on the HCP dataset. Specifically, we trained HMM and DyNeMo models with six states or modes on the HCP data, and extracted the corresponding state/mode time courses and covariance matrices. We then simulated new datasets by randomly selecting 500 subjects, extracting the first-session state/mode time courses (1200 time points), and generating synthetic time series by sampling independent multivariate Gaussian observations at each time point. For the HMM, each time point was drawn from the covariance matrix corresponding to the active state. For DyNeMo, we sampled from a Gaussian distribution with the covariance given by a weighted linear combination of all mode covariances, using the time-varying mixture weights as coefficients.

Because some DyNeMo modes learned from real data capture weak patterns, they can be difficult to recover in subsequent simulations, particularly when evaluated with a deep-learning model such as DyNeMo. To separate this data-related effect from the intrinsic behaviour of bi-cross-validation, we therefore performed an additional DyNeMo simulation in which both the mode time courses and mode covariances were randomly generated rather than derived from the trained model. This control analysis, described in Appendix B, served to verify that the conservative bias observed under bi-cross-validation in the realistic DyNeMo scenario arises from the uneven prominence of real-data modes rather than from limitations of the evaluation procedure.

Importantly, none of the simulations included temporal autocorrelation or convolution with the haemodynamic response function (HRF), as these effects are not yet incorporated into the current model frameworks. Extending the models to capture these temporal properties and generating corresponding simulations is an important direction for future work.

## 3 Results

### 3.1 Bi-cross-validation Recovers the Ground-Truth in Simulated Data

To evaluate the effectiveness of bi-cross-validation, we tested whether it could accurately recover groundtruth model structure in simulated datasets. Specifically, we simulated data from two models: an HMM with six states and a DyNeMo model with six modes (see Section 2.4 for details). We then applied bi-cross-validation to HMM, DyNeMo, and SWC, varying the number of estimated states or modes from 1 to 16, and compared their performance (Figure 3). For all three methods, log-likelihood values were demeaned relative to the same static baseline (*N*_states_ = 1), thereby making the y-axes directly comparable across panels (see Section 2.2.3).

**Figure 3.**
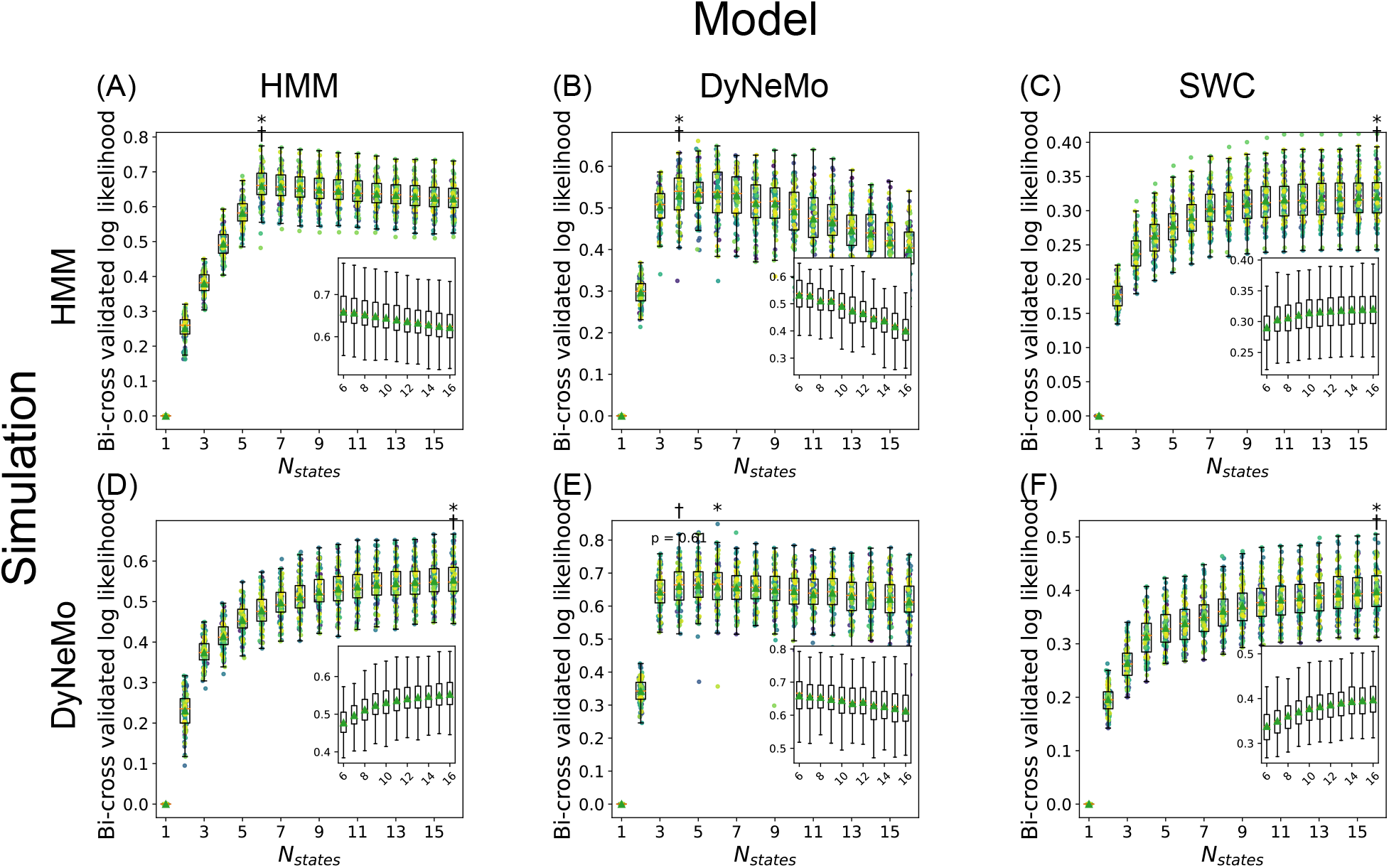
Simulation results. (A)–(C) Simulations based on the HMM generative model (6 states). (D)–(F) Simulations based on the DyNeMo generative model (6 modes). Columns correspond to evaluations of HMM, DyNeMo, and SWC using bi-cross-validation. In each panel, the x-axis indicates the number of states or modes used during model fitting, and the y-axis shows the bi-cross-validated log-likelihood. Each dot represents a single bi-cross-validation realisation, demeaned by the corresponding static model baseline (i.e., *N*_states_ = 1). Dots of the same color across values of *N*_states_ correspond to the same “reshuffle-and-split” realisation. Box plots show the distribution of log-likelihoods across realisations. An asterisk indicates the best-performing hyperparameter in each panel. A dagger marks the smallest number of states or modes whose performance is not significantly worse than the best-performing model, based on a one-sided paired t-test (*p <* 0.001). Insets show a zoomed-in view of the range *N*_states_ = 6 to 16 to highlight differences in model performance.

In the first simulation, where the data were generated from an HMM with six states, bi-cross-validation correctly identified the HMM with six states as the best-performing model, with a log-likelihood of 0.659 ± 0.049 (Figure 3A). Models with more than six states were penalised due to overfitting. DyNeMo underperformed relative to HMM, reaching its best performance at four modes (log-likelihood = 0.529 ± 0.058) before declining as model complexity increased (Figure 3B). In contrast, the performance of SWC improved with the number of states and peaked at 16 (log-likelihood = 0.320 ± 0.032), but its adjusted log-likelihood values (relative to the static baseline) remained substantially lower than those of HMM and DyNeMo (Figure 3C).

In the second simulation, we generated data from a DyNeMo model with six modes (Figure 3D–F). DyNeMo’s bi-cross-validated log-likelihood increased up to approximately four to six modes, where it reached a broad plateau (log-likelihood = 0.659 ± 0.070 at six modes), before declining with further increases in model order (Figure 3E), consistent with the partial recovery of ground-truth modes observed in the following sanity check analysis. HMM approached DyNeMo’s performance as the number of states increased, peaking at 16 states (log-likelihood = 0.553 ± 0.048), but remained suboptimal overall (Figure 3D). SWC consistently showed the weakest performance among the three methods, although it continued to improve with model complexity, reaching its best at 16 states (log-likelihood = 0.398±0.042; Figure 3F).

To better understand these results, we conducted a sanity check on the same simulated datasets. Specifically, we trained HMM and DyNeMo models on their respective full simulated datasets, using all subjects and all brain regions without any bi-cross-validation splits, under three settings: underestimating (*N*_states_ = 4), exact match (*N*_states_ = 6), and overestimating (*N*_states_ = 16). We then compared the recovered state or mode time courses and covariance matrices to the ground-truth (Figure 4 and Figure S1).

**Figure 4.**
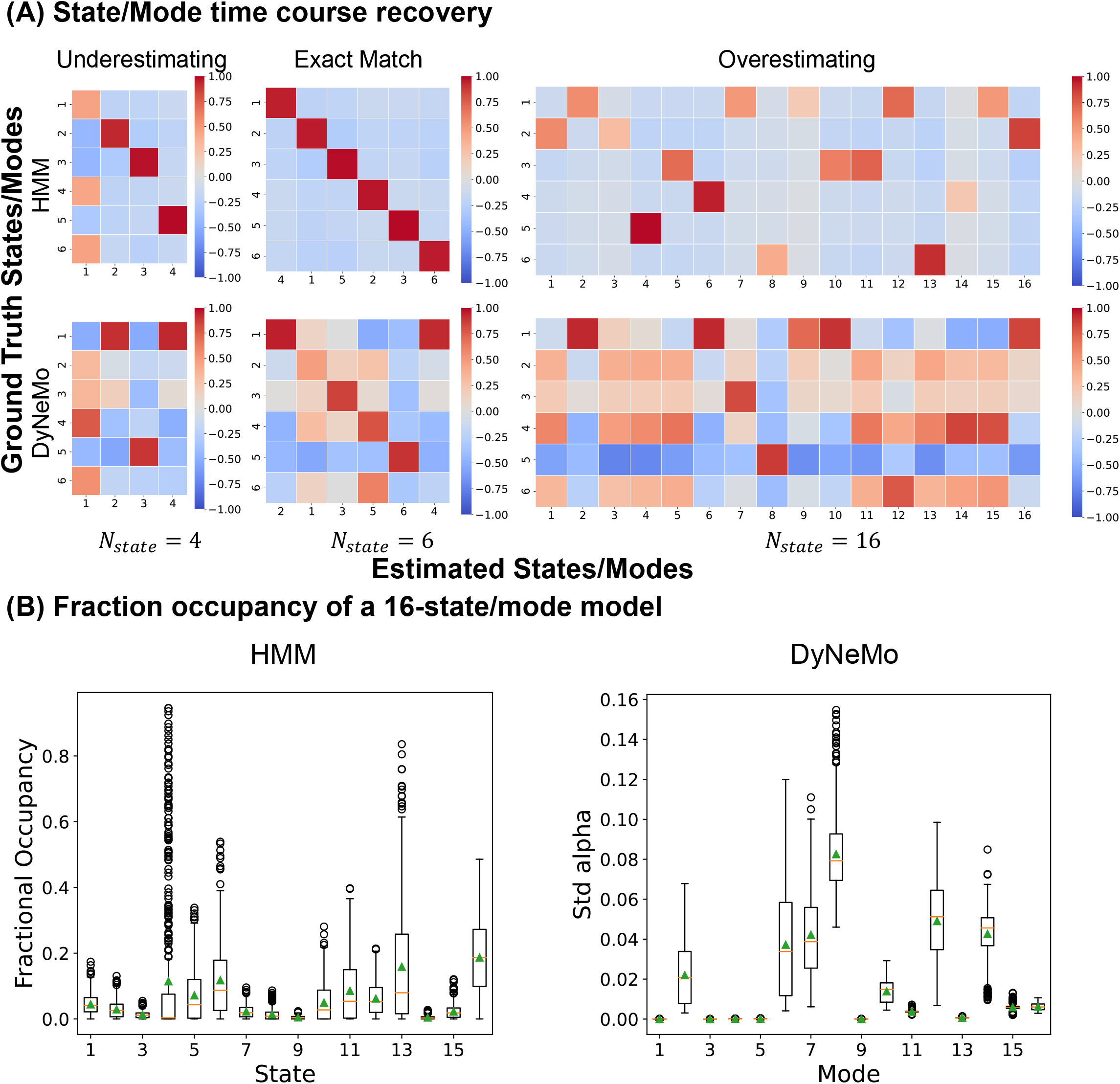
Temporal recovery analysis in HMM and DyNeMo simulations. (A) **State/mode time course recovery.** For the HMM 6-state simulation (first row), we trained three HMMs with 4 states (underestimating), 6 states (exact match), and 16 states (overestimating) on the full dataset. We then computed the pairwise correlation between the inferred and ground-truth state time courses. For the 6-state model, state correspondence was optimised using the Hungarian algorithm, attempting to optimally pair estimated to ground truth states. The second row shows the same procedure applied to the 6-mode DyNeMo simulation and DyNeMo estimation. (B) **Fractional occupancy distributions**. The distribution of fractional occupancy across simulated subjects for the overfitting 16-state HMM and 16-mode DyNeMo models.

In the HMM simulation, when the number of estimated states matched the ground-truth (*N*_states_ = 6), the recovered state time courses closely aligned with the true sequences (Figure 4A, first row, middle panel). When the number of states was too small (*N*_states_ = 4), the ground-truth time courses of only three states (states 2, 3, and 5) were recovered, while the remaining three were merged into a shared state (Figure 4A, first row, left panel). When the number of states was too large (*N*_states_ = 16), several states mapped imperfectly onto a single ground-truth state, leading to redundancy and fragmentation. For example, states 2, 7, 12, and 15 in the inferred model all corresponded to ground-truth state 1 (Figure 4A, first row, right panel). We also examined the fractional occupancy of each state (Figure 4B). Most states showed non-zero activation, even when some appeared to be noisy duplicates of the same underlying ground-truth state. Riemannian distances between estimated state covariances and ground-truth covariances (Figure S1) further confirmed that both underestimating and overestimating the number of states distort the temporal and spatial structure of the recovered states.

In the DyNeMo simulation, most ground-truth modes were recovered when the number of modes matched the ground-truth (*N*_modes_ = 6). Specifically, four modes were robustly recovered, while a fifth mode (mode 2) was only partially recovered and one mode was largely missed (Figure 4A, second row, middle panel). This incomplete recovery implies an effective dimensionality closer to four than six. This ambiguity is amplified under bi-cross-validation, where each evaluation uses only half of the brain regions and subjects. As a result, the bi-cross-validated log-likelihood plateaus after *N*_modes_ = 4 in Figure 3E. In this sense, bi-cross-validation behaves as a conservative estimator for DyNeMo. The underestimating and overestimating patterns observed in DyNeMo mirror those seen in HMM (Figure 4A, second row, left and right panels). However, DyNeMo tends to silence redundant modes rather than fragmenting existing ones, with many modes exhibiting near-zero activation (Figure 4B). A complementary analysis of mode covariance matrices is shown in Figure S1, further supporting these interpretations.

These observations highlight an intrinsic challenge arising from the interaction between DyNeMo and the simulated dataset. To confirm that the conservative bi-cross-validation pattern reflects this data-model interaction rather than a limitation of the evaluation procedure itself, we conducted an additional control simulation using synthetically generated mode time courses and covariance matrices (Appendix B). In this fully controlled setting, bi-cross-validation identified a near-correct model order of seven modes (Figure S2), indicating that the method is sensitive to model complexity but does not systematically underestimate it.

Together, these analyses explain the expected rise-then-fall pattern observed in the bi-cross-validated log-likelihood curves (Figure 3A and E): model performance improves as complexity approaches the ground-truth and declines once redundant states or modes are introduced. By jointly capturing spatial and temporal accuracy while penalising excessive complexity, bi-cross-validation provides a unified, data-driven metric for model evaluation. Across both HMM and DyNeMo simulations, it successfully recovered the underlying model structure and identified model orders consistent with the ground truth (or very close to it in the DyNeMo case). These results strengthen our confidence in applying bi-cross-validation to real data.

### 3.2 Bi-cross-validation for Model Selection in Real Data

Having validated bi-cross-validation using simulated data, we next applied the method to real restingstate fMRI datasets. Our primary analysis focused on data of 1003 subjects from the HCP young adult cohort (Smith et al., 2013), and we assessed the replicability of our findings using resting-state fMRI from 5000 randomly selected subjects in UKB (Miller et al., 2016). For both datasets, we analysed resting-state networks (RSNs) found using group-level ICA with 50 components (Smith et al., 2014). The effect of network dimensionality on model performance is explored in a later section.

The four candidate models — HMM, DyNeMo, SWC and static functional connectivity (sFC) — were introduced in Section 2.1. To ensure fair and robust comparisons, we used the same 100 “reshuffle-and-split” realisations across all models, varying the number of estimated states or modes from 2 to 25. For the static functional connectivity (sFC) model, the number of “states” corresponds to the number of clusters of subject-level static FC matrices across subjects, rather than temporally varying states within subjects. In each bi-cross-validation fold, the model combines its inferred state or mode time courses with spatial observation model parameters, to reconstruct a multivariate Gaussian distribution at each time point and computes the corresponding log-likelihood. The final scoring function is the bi-cross-validated log-likelihood, defined as the average log-likelihood per time point. We subtracted the log-likelihood of the *N*_states_ = 1 model under the same “reshuffle-and-split” realisation (see Section 2.2.3). Because this baseline corresponds to a common 1-state model across all four approaches and the splits were identical across subjects, the demeaned log-likelihood values are placed on the same scale, allowing direct comparison across models. We selected the number of states or modes using the “best-median-then-backward search” strategy described above.

The bi-cross-validation results for the HCP dataset (50-component ICA) are shown in Figure 5. For HMM, bi-cross-validation identified 9 states as the model with the highest median log-likelihood, but performance plateaued between 6 and 9 states, before declining with additional complexity (Figure 5A). Based on our criteria, the final selected model used 6 states, with a log-likelihood of 0.312 ± 0.044. For DyNeMo, although the highest median bi-cross-validated log-likelihood was observed at 25 modes (Figure 5B), model performance plateaued at 14 modes, yielding a log-likelihood of 0.633±0.101, with no significant improvement thereafter. SWC showed a monotonic increase in bi-cross-validated log-likelihood across the initial range of 2 to 25 states (Figure 5C), reaching 0.328 ± 0.036 at 24 states. To assess whether this trend reflected meaningful structure or continued over-partitioning, we extended the search range to 100 states in steps of 5. Within this extended range, performance peaked at approximately 50 states (Figure S3); however, the total improvement was minimal, amounting to less than 0.03 in log-likelihood despite a substantial increase in model complexity. For static functional connectivity (sFC), performance peaked around 10 clusters (log-likelihood = 0.312 ± 0.0389), followed by a minor but consistent downward trend beyond that point (Figure 5D). When comparing across models, DyNeMo achieved substantially higher bi-cross-validated log-likelihood under the common probabilistic framework, consistent with the benefits of relaxing the non-overlapping state assumption and allowing more flexible, long-range temporal dynamics. In contrast, HMM, SWC, and sFC exhibited similar peak log-likelihoods despite their differing temporal assumptions.

**Figure 5.**
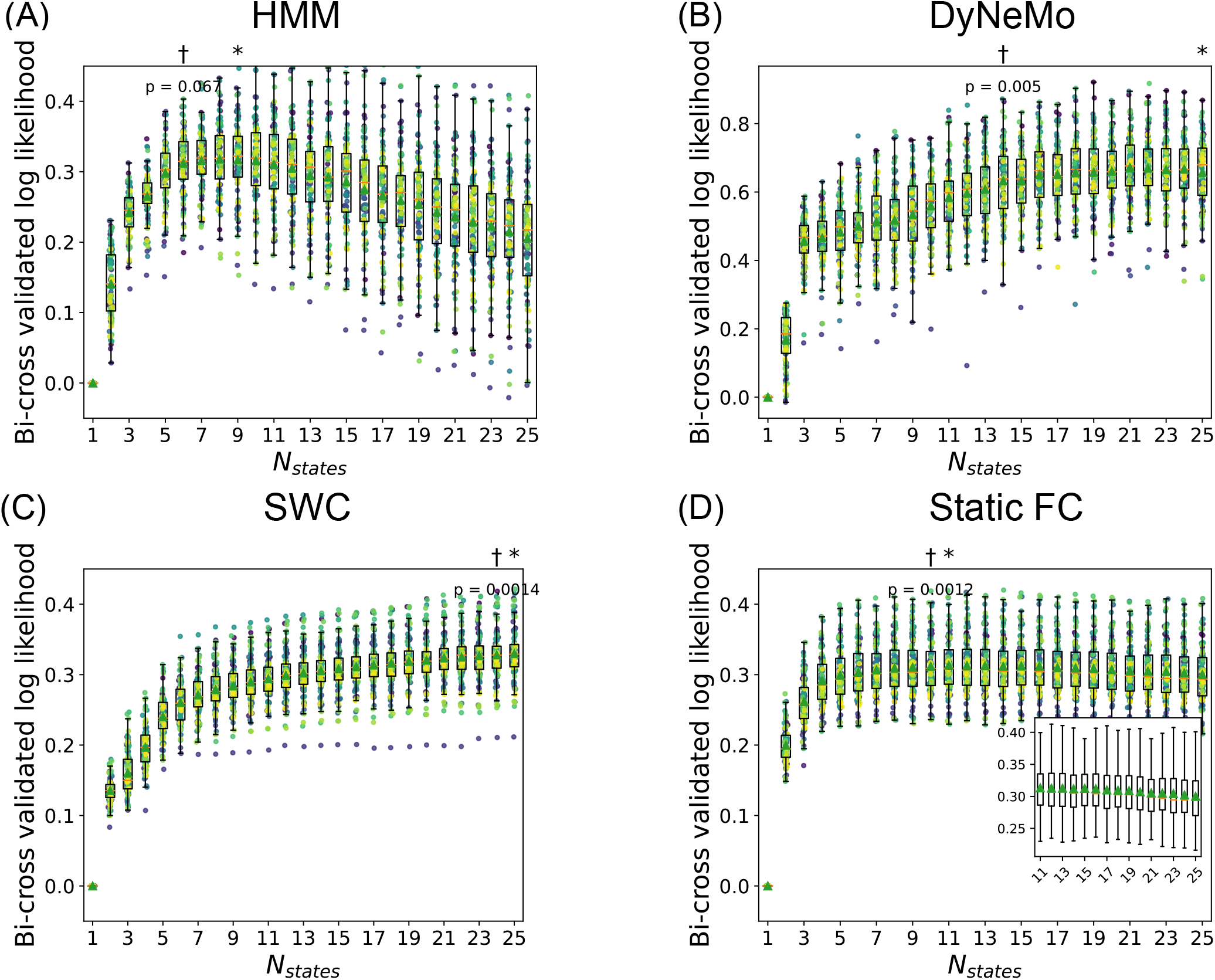
Model performance on HCP ICA 50. (A) HMM, (B) DyNeMo, (C) SWC (D) static FC. Resting-state fMRI data from HCP were processed with group-level spatial ICA with 50 components, followed by dual regression to obtain subject-level time series. Bi-cross-validation was applied to each model with number of states ranging from 1 to 25. Each dot represents a bi-cross-validation realisation; boxplots summarise the distribution across 100 reshuffle-and-split repetitions. An asterisk indicates the model order with the highest median bi-cross-validated log-likelihood. A dagger marks the smallest number of states or modes whose performance is not significantly worse than the best-performing model, based on a one-sided paired t-test (*p <* 0.001). The inset in panel (D) provides a zoomed-in view over *N*_states_ = 11 to 25 to highlight differences in model performance.

To assess the replicability and robustness of our findings, we applied bi-cross-validation to the UKB dataset using the same 50-component ICA decomposition (Figure S4). For HMM, performance plateaued between 5 and 11 states; under our selection criterion, the final model used 5 states (log-likelihood = 0.371 ± 0.0619; Figure S4A). Bi-cross-validation selected 12 modes for DyNeMo (log-likelihood = 0.939±0.121; Figure S4B) and 23 states for SWC (log-likelihood = 0.309±0.0305; Figure S4C). For static functional connectivity (sFC), performance peaked at 12 clusters with log-likelihood = 0.400 ± 0.0444 (Figure S4D). Both within-model and across-model performance trends were largely consistent with those observed in the HCP data, with only minor differences in the optimal number of states or modes. These discrepancies may be attributed to the larger sample size and greater subject variability in UKB, as well as the shorter scanning sessions (490 time points in UKB vs. 4 sessions totalling 4800 time points in HCP).

We also examined the potential influence of low-frequency signal components on model evaluation, particularly for SWC, where previous studies have reported spurious correlations arising from interactions between slow fluctuations and the sliding-window procedure (Leonardi & Van De Ville, 2015). To probe this, we applied an aggressive fifth-order Butterworth high-pass filter (cut-off frequency at 0.05 Hz, corresponding to fluctuations slower than ~20 s) to the HCP dataset and repeated the bi-cross-validation analysis (Figure S5). Across models, the overall shape of the bi-cross-validated log-likelihood curves and the relative ordering of model performance were preserved after filtering, indicating that our main conclusions are not driven by low-frequency confounds. Notably, DyNeMo exhibited a clearer decline in performance at high model orders in the filtered data, suggesting that bi-cross-validation more strongly penalises overfitting once slow signal components are suppressed.

Finally, we sought to understand what additional information bi-cross-validation provides relative to conventional model evaluation metrics. In generative models such as HMM and DyNeMo, the free energy serves as the cost function during training and is commonly used as a measure of model fit. We therefore implemented a standard five-fold cross-validation procedure in which subjects were split across folds. In each fold, the model was trained on the training set, state or mode time courses were inferred for the test set, and the final free energy was computed. In addition, split-half reproducibility is frequently used in dFC modelling as an index of robustness. We quantified this using the Riemannian distance between state or mode covariance matrices estimated from two data halves (see Section 2.3 for details).

The comparison between bi-cross-validated log-likelihood, standard cross-validated free energy and split-half Riemannian distance is illustrated in Figure 6. For HMM, free energy continued to decrease monotonically with increasing number of states (Figure 6B), reflecting a known limitation of standard cross-validation, which favours increasingly complex models without identifying an optimal number of states (discussed in Section 2.2). Split-half reproducibility declined with increasing model complexity, as indicated by rising Riemannian distances between paired states estimated from the two data halves (Figure 6C). Notably, sharp decreases occurred when moving from 4 to 5 states for HMM and from 3 to 4 modes for DyNeMo, consistent with the general tendency of added complexity to reduce stability across data splits. As expected, the 1-state model showed the highest reproducibility, reflecting its minimal model complexity. In contrast, bi-cross-validated log-likelihood displayed an “increase-then-decrease” trend (Figure 6A), illustrating its ability to balance model fit and complexity. For DyNeMo, the pattern differed from HMM. As model complexity increases, free energy decreased (indicating improved fit) up to 12 modes, while split-half reproducibility simultaneously worsened, reaching its lowest point at 12 modes (Figures 6E and F). Beyond this point, free energy plateaued and reproducibility began to improve. This behaviour stems from DyNeMo’s model structure: when additional modes fail to explain variance, the model effectively silences them and sets their mode covariances toward the prior, which are more reproducible by design. We further explore and confirm this behaviour in the next section.

**Figure 6.**
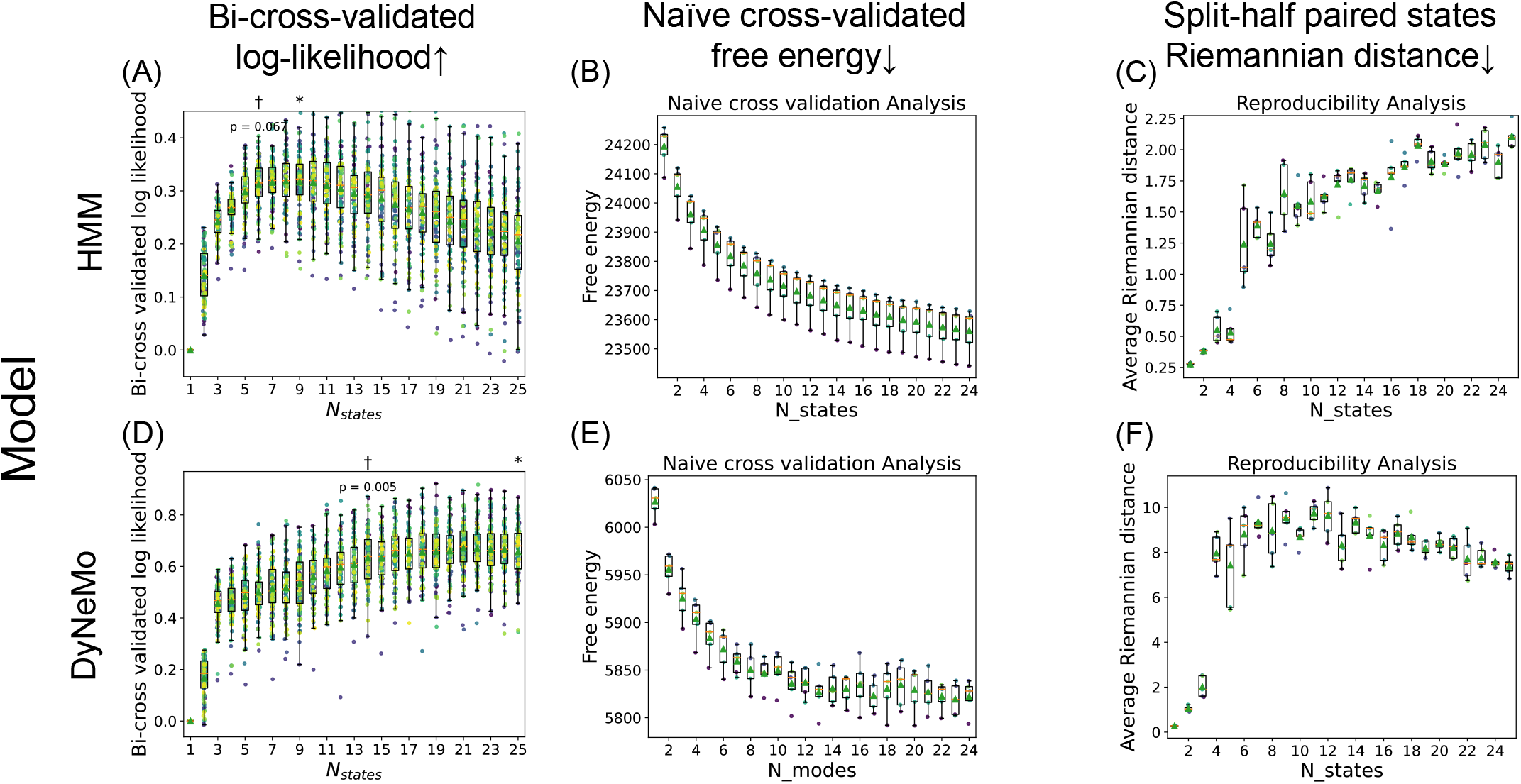
Comparison between bi-cross-validation and traditional metrics. (A)–(C) show results for the Hidden Markov Model (HMM), while (D)–(F) correspond to DyNeMo. All analyses were performed using the HCP data with a 50-component group-level ICA decomposition. Bi-cross-validation results (left column) are identical to those presented in Figure 5, where higher values indicate better performance (↑). The middle column shows results for standard five-fold cross-validated free energy, where lower values indicate better performance (↓). The right column shows split-half reproducibility, quantified as the average Riemannian distance between paired state or mode covariances across data halves; here again, lower values indicate better reproducibility (↓). In free energy and reproducibility analyses, models were trained separately on each split or fold, and comparisons were made as described in Section 2.3.

These results highlight the effectiveness of bi-cross-validation for evaluating dFC models. By implicitly balancing model fit and complexity, it offers a principled framework for both within-model and across-model selection, as well as robust comparison of dynamic models against static functional connectivity baselines.

### 3.3 Post-Hoc Validation: Temporal and Spatial Interpretability

Having examined bi-cross-validation results on real data (Figure 5), we conducted additional post-hoc analyses to explain the observed trends and further validate our findings.

We first address the following question: why does the bi-cross-validated log-likelihood on real data decline sharply with increasing number of states in HMM (Figure 5A), but plateau in DyNeMo (Figure 5B)? To investigate this, we fitted both an HMM and DyNeMo with 25 states or modes to the full dataset and examined the fractional occupancy of each (see Section 2.3.3 for details; Figure 7). In HMM, the mean fractional occupancy was non-zero for nearly all 25 states (Figure 7B), indicating that each state was visited for a measurable proportion of time across subjects. In contrast, DyNeMo exhibited a different pattern: 11 modes were effectively “silenced” (Figure 7E), consistent with the behaviour observed in simulations (Figure 4B). This closely mirrors the bi-cross-validation results, where DyNeMo performance plateaued after 14 modes. To further characterise these differences, we ranked states and modes by their fractional occupancy and examined the pairwise Riemannian distances between their corresponding covariance matrices. In HMM, highly occupied states were relatively similar to one another, while rare states were increasingly distinct (Figure 7C). In DyNeMo, modes with low fractional occupancy clustered near the prior covariance (Figure S6), indicating that these modes contributed little additional structure. In contrast, DyNeMo modes with high fractional occupancy exhibited larger pairwise Riemannian distances than HMM states (Figure 7F), consistent with DyNeMo modes capturing dominant directions of covariance variation rather than discrete brain states.

**Figure 7.**
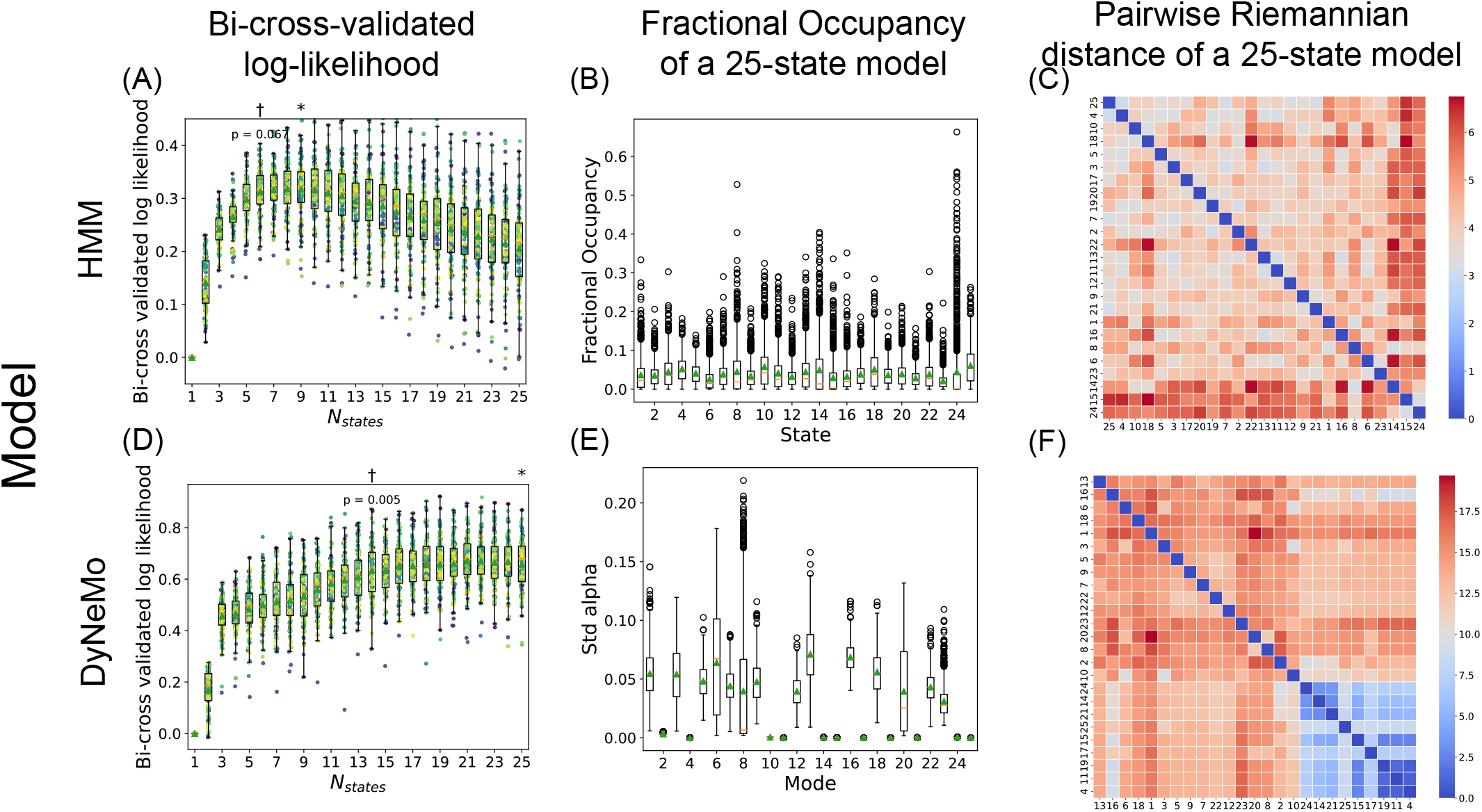
Post-hoc Analysis on HMM and DyNeMo (HCP ICA-50). (A,D) Bi-cross-validated log-likelihood as a function of model order for HMM (A) and DyNeMo (D), reproduced from Figure 5A–B for reference. (B,E) Fractional occupancy profiles for a 25-state HMM (B) and a 25-mode DyNeMo model (E). For HMM, fractional occupancy is computed as the proportion of time each subject spends in each state, with distributions shown across subjects. For DyNeMo, an analogous measure of mode prominence is defined as the standard deviation of the posterior mode time courses, normalised by the corresponding mode covariances (Section 2.3.3). (C,F) Pairwise Riemannian distances between HMM state (C) or DyNeMo mode (F) covariance matrices, ordered by decreasing fractional occupancy.

These observations suggest a difference in how model complexity is expressed in the two approaches: HMM incrementally adds rare and spatially distinct states as complexity increases, which may be less reproducible and therefore more heavily penalised by bi-cross-validation. DyNeMo, on the other hand, captures the dominant spatiotemporal patterns early and silences any redundant modes by setting their covariances close to the prior. As a result, this mechanism appears to suppress overfitting, and unused modes contribute minimal complexity. These silent modes may capture subtle, low-frequency structure in the data, potentially contributing to slight performance gains in unfiltered data (Figure 5B) and corresponding drops after high-pass filtering (Figure S5B).

We next examined whether the model orders recommended by bi-cross-validation correspond to meaningful spatial structure in the data. As shown in Figure 5, a six-state HMM and a 14-mode DyNeMo achieved near-optimal performance under bi-cross-validation. To assess the interpretability of these solutions, we trained an HMM with six states and a DyNeMo with 14 modes on the HCP ICA 50 dataset and visualised their state- or mode-specific spatial patterns (Figure 8; full DyNeMo results in Figure S7). For each state or mode, we extracted the covariance matrix, converted it to a correlation matrix, and computed a rank-one approximation by extracting the dominant eigenvector. This eigenvector was then projected back into brain space to obtain a spatial map for visualisation. To characterise the spectral structure underlying this approximation, we also examined the eigenspectra of the state and mode correlation matrices (Figure S8).

**Figure 8.**
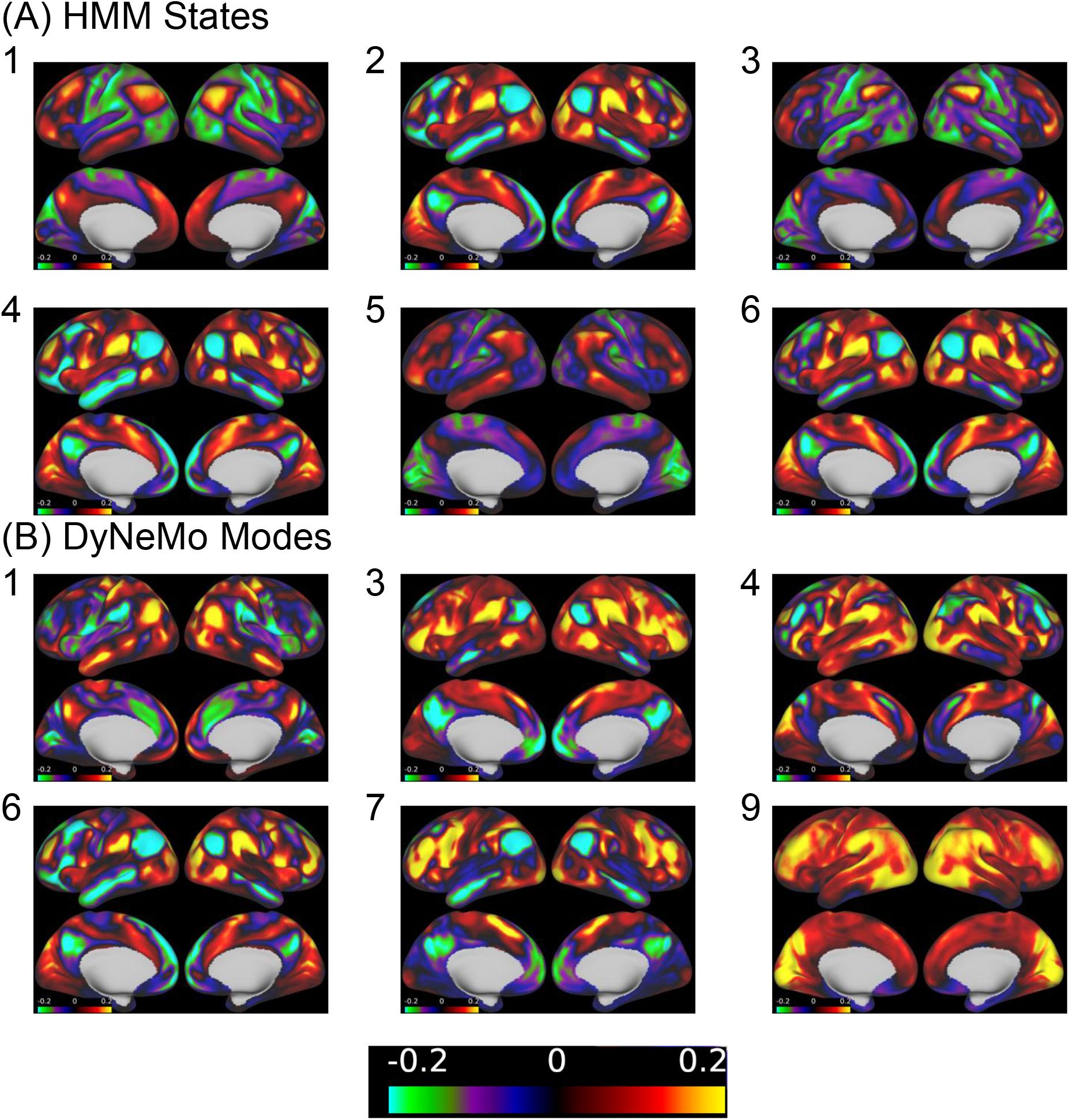
Visualisation of HMM states and DyNeMo modes (HCP ICA 50). Spatial maps derived from a six-state HMM (A) and a 14-mode DyNeMo (B) trained on the HCP ICA 50 dataset. For both models, state or mode covariance matrices were extracted after training and converted to correlation matrices. For each state or mode, a rank-one approximation was computed by extracting the dominant eigenvector (i.e., the eigenvector associated with the largest eigenvalue) of its correlation matrix. This eigenvector was projected back to brain space to obtain a spatial map for visualisation. All six HMM states are shown. For DyNeMo, six representative modes are displayed in the main figure; the complete set of 14 modes is provided in Figure S7. A single symmetric colour scale centred at zero was used for all states and modes.

The six HMM states correspond broadly to well-known large-scale resting-state networks (Figure 8A). For example, State 1 primarily reflects the default mode network (DMN), State 2 emphasises the ventral attention network, and States 3 and 5 resemble components of the frontoparietal network. These patterns are broadly consistent with canonical parcellations such as Yeo et al.’s 7-network solution (Thomas Yeo et al., 2011). However, we also observe partial redundancy among states. In particular, States 2, 4 and 6 all prominently involve the ventral attention network, differing mainly in their spatial weighting. One possible explanation is that the HMM is a group-level model that assumes a shared set of discrete states across subjects. Inter-subject variability in spatial organisation may therefore be partially expressed in the temporal domain, leading to multiple states that reflect similar large-scale networks but differ in how they are expressed across individuals. In this sense, some of the apparent state diversity may reflect inter-subject variability rather than distinct functional configurations. The eigenspectra of the HMM state correlation matrices (Figure S8A) show that the ratio between the largest eigenvalue and the sum of all eigenvalues ranged from below 0.15 in three states to approximately 0.25 in the most dominant case, indicating that the leading eigenvector captured only a modest proportion of the total spectral mass. The corresponding first eigenvectors were relatively diffuse across ICA components, with cumulative energy distributed over many components rather than concentrated in a small subset.

In contrast, the 14-mode DyNeMo solution exhibits a more compositional structure. While some modes closely resemble canonical networks (e.g., Mode 1 corresponding to the DMN, Mode 3 to the ventral attention network, and Mode 9 to the visual network), others capture coordinated patterns spanning multiple large-scale systems. For example:

1. Mode 4 involves both visual and ventral attention networks.
2. Mode 6 combines ventral attention and frontoparietal components.
3. Mode 7 links frontoparietal and dorsal attention networks.

These modes are therefore not restricted to single canonical systems but instead reflect coordinated interactions between networks. When compared with finer-grained parcellations (e.g., Yeo et al.’s 17-network), additional structure becomes apparent. For instance, although Modes 6 and 7 both involve frontoparietal networks, Mode 6 aligns more closely with Network 8 in Yeo-17, whereas Mode 7 more closely resembles Network 12. Similarly, while Modes 1 and 2 both map onto the DMN, Mode 1 appears more closely related to a DMN subnetwork (Network 16 in Yeo-17).

As with the HMM, some degree of repetition is present (e.g., Modes 2 and 10, Modes 6 and 13), which may again reflect subject-level variability or limitations of the group-level ICA decomposition. Nevertheless, the DyNeMo solution more frequently captures structured combinations of networks rather than slight variants of a single system. This compositional structure is further reflected in the eigenspectra of the DyNeMo mode correlation matrices (Figure S8B), where the largest eigenvalue ratio ranged from approximately 0.10 to 0.50, with several modes exceeding the spectral concentration observed in any HMM state. This suggests that, for a subset of modes, connectivity patterns are concentrated along a dominant low-dimensional direction. Together, these spatial and spectral differences are consistent with the superior bi-cross-validation performance observed for DyNeMo (Figure 5), indicating that its more compositional representation may better capture organised network dynamics.

### 3.4 ICA Dimensionality Affects Model Performance

ICA provides a flexible framework for estimating RSNs at varying levels of dimensionality, by adjusting the number of extracted components. Leveraging this flexibility, we evaluated how ICA dimensionality affects dFC model performance. While earlier sections focused on 50-component ICA, we extended our analysis to include ICA with 15, 25 and 100 RSNs. Higher-dimensional RSN decompositions were not examined due to computational constraints.

Model comparison results for the 15-component ICA are shown in Figure 9. At this lower dimensionality, both HMM and DyNeMo performed worse than the static single-state baselines (Figure 9A and B). Although SWC showed an improvement over the static single-state baseline at low model orders (Figure 9C), its performance declined as the number of states increased, and it remained inferior to the static functional connectivity model obtained by clustering sFC across subjects (Figure 9D). A similar pattern was observed for the 25-component ICA (Figure S9). In contrast, the 100-component results closely resembled those obtained with 50 components, with dynamic models clearly outperforming their static counterparts (Figure S10). Notably, while HMM performed similarly to sFC and SWC at 50 components (Figure 5A), at 100 components it achieved substantially higher log-likelihood than both static and sliding-window approaches, ranking second only to DyNeMo. DyNeMo remained the best-performing model, reinforcing the advantage of flexible dynamic formulations at higher spatial resolutions. To assess the generalisability of our findings, we also applied the 25-component ICA to the UKB dataset. The overall pattern was consistent with the 25-component HCP results, with static models outperforming most dynamic approaches, although DyNeMo showed relatively stronger performance in this dataset (Figure S11B).

**Figure 9.**
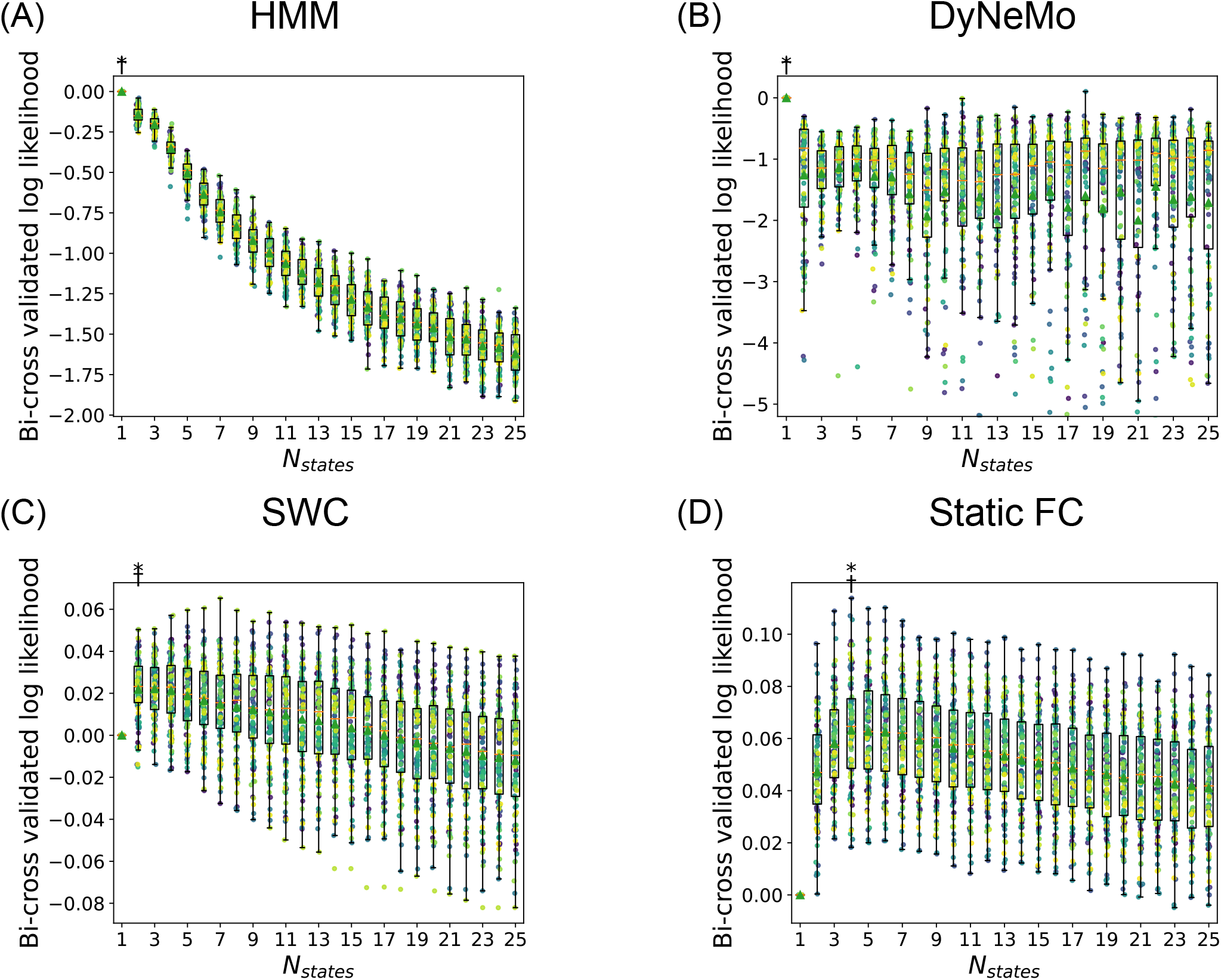
Model performance on HCP ICA 15. (A) HMM, (B) DyNeMo, (C) SWC (D) static FC. Resting-state fMRI data from the HCP were processed using group-level spatial ICA with 15 components, followed by dual regression to extract time series. Bi-cross-validation was applied to each model across model orders ranging from 1 to 25. All plots were generated using the same procedure as in Figure 5 with asterisks marking the model with the highest median log-likelihood, and daggers (†) indicating the smallest model not significantly worse than the best (one-sided paired t-test, *p <* 0.001). In this case, the dagger coincides with the asterisk, indicating that reducing the model order by even one leads to a significant drop in performance.

A general framework for interpreting these results is that dynamic functional connectivity effects become more pronounced with increasing ICA dimensionality. As we move from low-dimensional ICA (15 and 25 regions) to higher-dimensional ones (50 and 100 regions), dFC model performance gradually improves. There may exist a threshold or “turning point” in granularity beyond which dynamic models begin to outperform static models. The location of this turning point may vary across datasets.

## 4 Discussion

In this paper, we introduced bi-cross-validation as a data-driven framework for evaluating dFC models. This approach enables fair and robust comparisons across models with different hyperparameter settings, provided they offer a whole-brain, spatiotemporal probabilistic representation of neuroimaging data. The core assumption underlying bi-cross-validation is that large-scale brain network dynamics involve multiple regions. To adhere to this assumption as closely as possible, we implemented a multi-instance, two-fold “reshuffle-and-split” strategy across both subjects and brain regions.

We validated the method using simulated data, demonstrating its ability to recover the ground-truth model. We then applied bi-cross-validation to resting-state fMRI data, where it consistently recommended model configurations that balanced goodness-of-fit and complexity. Finally, we investigated the influence of ICA dimensionality on dFC modelling, showing that finer-grained ICAs enhance dFC model performance as assessed by bi-cross-validation, whereas coarser representations tend to obscure dynamic effects. Taken together, these findings address three central questions in dFC modelling:

1. Does functional connectivity exhibit meaningful temporal dynamics (i.e., dynamic vs. static FC) (Laumann et al., 2017)
2. How should the key hyperparameters be selected during model fitting (within-model evaluation)? (Allen et al., 2014; Gohil et al., 2022; Vidaurre et al., 2017)
3. How can different dFC models be compared objectively and consistently (across-model evaluation)? (Lurie et al., 2020)

### 4.1 Bi-cross-validation guides further model development

Bi-cross-validation provides valuable guidance for neuroimaging methodologists during model development. A crucial aspect of model fitting is selecting appropriate hyperparameter settings (Allen et al., 2014; Gohil et al., 2022; Vidaurre et al., 2017). We have demonstrated bi-cross-validation’s effectiveness in determining the optimal number of states or modes, but this technique can also be extended to fine-tune other hyperparameters. For example, researchers employing sliding window correlation (SWC) could use bi-cross-validation to optimise window length, window type, and clustering method (Shakil et al., 2016).

Additionally, bi-cross-validation aids in making informed decisions about model structure. DyNeMo benefits from its more flexible formulation, which relaxes both the mutually exclusive state assumption and the Markovian temporal constraint of the HMM (Gohil et al., 2022). Similar analyses can be used to evaluate whether incorporating haemodynamic response functions (Lindquist et al., 2009), mean activation (Quinn et al., 2018), or subject-embeddings (Huang et al., 2026) provides substantial improvements. More importantly, bi-cross-validation offers a fair framework for inter-model comparisons, enabling researchers to demonstrate the strengths of their models. Our implementation is designed to be general and user-friendly, facilitating the testing of novel models.

### 4.2 Bi-cross-validation supports dFC model applications

Bi-cross-validation is also valuable for researchers applying dFC models to their own datasets. A common challenge faced by experimental neuroimaging researchers is selecting appropriate model hyperparameters, particularly the number of states. Bi-cross-validation can assist in this process by systematically evaluating models and hyperparameters on preprocessed large-scale resting-state fMRI datasets.

We acknowledge that model performance is influenced by the preprocessing pipeline, parcellation method, and dataset size. Consequently, we encourage researchers to ensure proper data preprocessing, select an appropriate parcellation, and compare multiple models using bi-cross-validation before drawing conclusions from their results.

### 4.3 The role of ICA dimensionality in dFC modelling

One notable finding of this study is that the performance of dFC models depends strongly on ICA dimensionality. At lower dimensionalities, static models provided a better description of the data, whereas at higher dimensionalities, dynamic models became increasingly advantageous. Notably, the advantage of discrete-state models such as the HMM emerged only at the highest spatial resolution. This aspect has been largely overlooked in the literature. For instance, Laumann et al. (2017) primarily focused on a dense parcellation. Similarly, many impactful dFC studies (Allen et al., 2014; Vidaurre et al., 2017) employ a fixed parcellation scheme without systematically assessing how parcellation granularity affects model performance. Future model development and evaluation should incorporate comparisons across different parcellation schemes to better understand their influence.

This finding also has implications for understanding brain organisation. As parcellations shift from coarse to fine-grained networks, large-scale neural systems are often subdivided into subnetworks (Schaefer et al., 2018). Our results suggest that dynamic interactions occur between these subnetworks (Farahibozorg et al., 2025), but when their signals are merged into broader networks and averaged, these dynamics may be blurred or diminished. Investigating the hierarchical organisation of brain dynamics and how dynamic interactions manifest across different spatial scales represents an important direction for future research.

### 4.4 Limitations

Despite its advantages, bi-cross-validation has several practical limitations. Here, we outline key challenges and potential strategies to address them. First, bi-cross-validation is computationally demanding, particularly when applied to large datasets. In this study, we performed 100 reshuffle-and-split iterations to assess model performance robustly. However, users may achieve reasonable estimates with 10–20 iterations, or increase the number if needed. Second, bi-cross-validation cannot inherently distinguish between meaningful neural dynamics and systematic noise. As a data-driven approach, it focuses on how well a model describes the data. If a model consistently captures artefacts such as scanner drift, head motion or physiological noise (Laumann et al., 2017; Power et al., 2017), bi-cross-validation may not penalise this. Nevertheless, our results indicate that the dynamic effects observed in the 50- and 100-component ICAs are unlikely to be purely driven by global noise. If artefacts such as head motion or physiological signals were the main drivers of dFC states, similar effects should be present across all ICA dimensionalities. Instead, we found that models with 15- and 25-component ICAs showed little evidence of dynamics, whereas finer-grained ICAs revealed robust patterns. Importantly, these results were stable when applying additional high-pass filtering, further suggesting that the observed effects are not simply explained by low-frequency drifts. Moreover, while we used high-quality HCP data (Smith et al., 2013) for our primary analyses, we also examined UKB data, characterised by greater noise levels, to assess replicability. To mitigate the impact of artefacts, users should ensure that rigorous preprocessing pipelines are applied before using bi-cross-validation for model selection. Third, bi-cross-validation does not directly provide guidance on selecting imaging-derived phenotypes for downstream analyses. A common approach in resting-state fMRI research is to extract phenotypes from dFC models for use in behavioural or clinical prediction tasks (Damaraju et al., 2014; Griffin et al., 2024). While bi-cross-validation identifies the model that best describes the fMRI data, it does not indicate whether this model will produce features that are optimal for prediction. We argue that accurate representation of the underlying fMRI data is a necessary but not sufficient condition for generating predictive imaging-derived phenotypes.

We acknowledge that several dFC models fall outside the scope of our current evaluation framework. For example, co-activation pattern analysis (CAP) (Liu & Duyn, 2013; Liu et al., 2018) identifies recurring activation patterns from a subset of frames while discarding the rest, which complicates direct comparisons with other models. However, since CAP also relies on clustering to summarise recurrent patterns, bi-cross-validation could still be used to optimise the number of centroids. Additionally, some methods analyse functional connectivity dynamics in the frequency domain (Chang & Glover, 2010; Yaesoubi et al., 2015), which cannot yet be evaluated within our current framework. We aim to generalise our evaluation method to include such models in future work.

### 4.5 Future Work

Building on bi-cross-validation as a robust framework for evaluating dFC models, several avenues remain for future research. First, bi-cross-validation’s reliability should be further assessed across diverse datasets varying in acquisition protocols, preprocessing pipelines, parcellation schemes, and sample sizes. Model performance is influenced by these factors, and sample size requirements differ across models. For example, SWC and HMM may perform adequately on smaller datasets, whereas deep-learning-based models like DyNeMo typically require larger samples. Second, bi-cross-validation can be extended to evaluate dFC models in other neuroimaging modalities, particularly MEG and EEG, where brain dynamics unfold at faster timescales. Applying bi-cross-validation to these modalities would not only strengthen confidence in dFC modelling, but also help validate its utility across functional neuroimaging techniques. Finally, future work will focus on using bi-cross-validation to guide the development of new dFC models. This includes building models that incorporate features such as autocorrelation structure, haemodynamic response modelling, or mean activation differences across states, with the goal of achieving a more comprehensive characterisation of dynamic changes in large-scale brain networks.

## Ethics statement

This study uses publicly available data from the Human Connectome Project (HCP) and UK Biobank (UKB). The HCP study protocol was approved by the Washington University–Minnesota Consortium institutional review boards, and written informed consent was obtained from all participants prior to data collection. UK Biobank has ethical approval from the North West Multi-Centre Research Ethics Committee (MREC) to obtain and disseminate data and samples from participants, and all participants provided informed consent. These ethical approvals cover the use of UK Biobank data in the present study. No additional ethical approval was required for this secondary analysis of existing data.

## Data and Code Availability

Human Connectome Project (HCP) data are available upon registration via the HCP website.

UK Biobank data were used under Application Number 8107 and are available upon registration and application for data access via the UK Biobank website.

The analysis code implementing the bi-cross-validation framework and associated modelling procedures is available at https://github.com/weiym97/osl-dynamics/tree/WIP, and will be incorporated into the main `osl-dynamics` repository (https://github.com/OHBA-analysis/osl-dynamics) upon publication.

## Author Contributions

Yiming Wei: Conceptualisation; Methodology; Software; Formal analysis; Investigation; Data curation; Visualisation; Writing – original draft.

Stephen M. Smith: Conceptualisation; Data curation; Funding acquisition; Methodology; Investigation; Resources; Supervision.

Chetan Gohil: Software.

Rukuang Huang: Conceptualisation; Methodology.

Ben Griffin: Methodology; Validation.

SungJun Cho: Software; Validation; Visualisation.

Stanislaw Adaszewski: Supervision.

Stefan Frässle: Supervision.

Mark W. Woolrich: Conceptualisation; Methodology; Investigation; Supervision; Funding acquisition.

Seyedeh-Rezvan Farahibozorg: Conceptualisation; Methodology; Investigation; Supervision; Funding acquisition.

All authors contributed to reviewing and editing the manuscript.

## Use of Generative AI

The manuscript was edited with the assistance of ChatGPT (OpenAI) to improve clarity and language. AI tools were also used to assist with drafting portions of analysis code. ChatGPT was accessed through the University of Oxford’s institutional provision (ChatGPT Edu). All outputs were carefully reviewed, unit-tested, and validated by the authors. All scientific conclusions and interpretations were developed and verified by the authors, and key findings were independently reproduced.

## Acknowledgements

We are grateful to the Human Connectome Project and UK Biobank for making these invaluable resources available, and to all participants who contributed their time to make these data possible. We thank members of the FMRIB Analysis Group and the OHBA Analysis Group for helpful discussions and constructive feedback during the development of this work. We thank Dr. Fidel Alfaro-Almagro for providing the UK Biobank ICA time series used in this study.

## Supplementary Material

The supplementary material is included in this PDF file after

## A Mathematical Details of Bi-cross-validation

In this section, we provide a detailed mathematical description of the bi-cross-validation framework introduced in Section 2.2. For clarity, we separately outline the generative model, the inference procedure, and the four-step bi-cross-validation evaluation for each class of models: Hidden Markov Model (HMM) (Vidaurre et al., 2017), Dynamic Network Modes (DyNeMo) (Gohil et al., 2022), and sliding window correlation (SWC) (Allen et al., 2014).

### Data and splitting conventions

Let there be *S* subjects, with subject *s* providing a time-by-region matrix 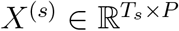, We construct

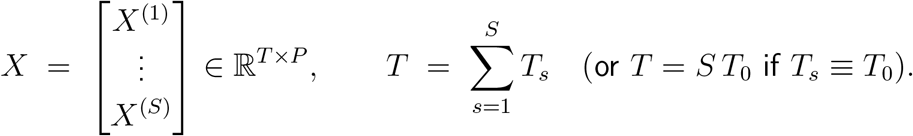

Bi-cross-validation partitions *rows* by *subjects* (never within a subject) and *columns* by regions. Let ℛ_train_, ℛ_test_ ⊂ {1, …, *T*} denote the row indices corresponding to a split of the subjects into two disjoint sets 𝒮_train_ and 𝒮_test_:

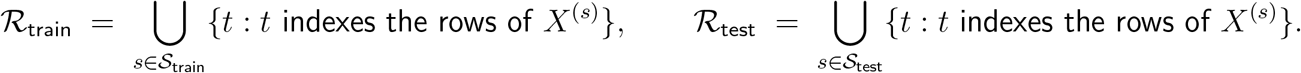

All time points for a subject (and all of that subject’s scanning sessions, if multiple are present) are assigned together to either ℛ_train_ or ℛ_test_. Within each subject or session, the temporal order of rows is preserved, reflecting the strong autocorrelation structure of fMRI time series.

Let 𝒞_*X*_, 𝒞_*Y*_ ⊂ {1, …, *P*} denote a column-partition of regions. The four bi-cross-validation quadrants are then

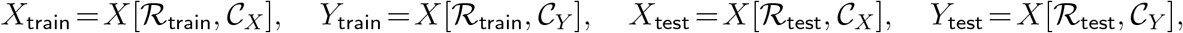

so that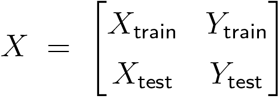, and final evaluation is performed on *Y*_test_ only.

### A.1 Hidden Markov Model (HMM)

#### A.1.1 Generative model

The HMM assumes that brain activity alternates between *K* discrete states. Let *s*_*t*_ ∈ {1, …, *K*} denote the latent state at time *t*, and **x**_*t*_ the observed fMRI vector. The joint distribution factorises as

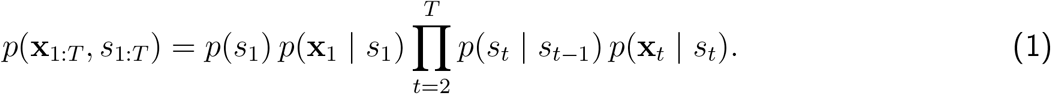

Each state *k* is associated with a multivariate Gaussian observation model,

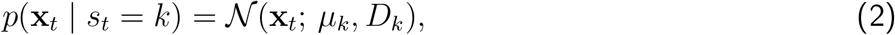

with mean vector *µ*_*k*_ and covariance matrix *D*_*k*_. Transitions are governed by a Markov transition matrix *A* = [*a*_*ij*_], where *a*_*ij*_ = *p*(*s*_*t*_ = *j* | *s*_*t*−1_ = *i*) and 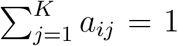. The distribution of the initial state is specified by *π* = (*π*_1_, …, *π*_*K*_), where *π*_*k*_ = *p*(*s*_1_ = *k*) and 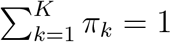.

#### A.1.2 Inference

Inference in `osl-dynamics` (Gohil et al., 2023) is based on a variational Bayesian formulation that minimises the variational free energy

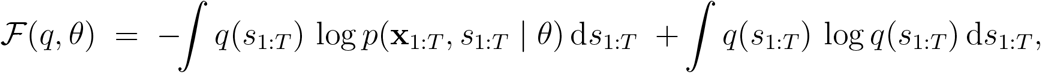

where *θ* = {*µ*_*k*_, *D*_*k*_, *A, π*} are the model parameters.

The posterior distribution over states is approximated by a factorised form

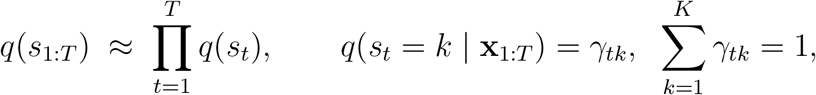

so that *γ*_*tk*_ gives the posterior probability of being in state *k* at time *t* given the observed data.

Optimisation proceeds by iteratively updating (i) the state probabilities *γ*_*tk*_, (ii) the observation parameters {*µ*_*k*_, *D*_*k*_}, and the Markov parameters (*A, π*), to reduce ℱ.

In summary, the inference scheme provides a probabilistic description of the latent state time courses through *γ*_*tk*_, while yielding point estimates for the observation model parameters *µ*_*k*_, *D*_*k*_ and for the transition parameters (*A, π*).

#### A.1.3 Bi-cross-validation procedure

Given a bi-cross-validation split, model evaluation proceeds as follows:

1. **Step 1 (Full train on** *Y*_**train**_): Estimate posterior time courses 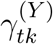 and parameters 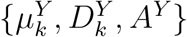 on *Y* _train_.
2. **Step 2 (Spatial inference on** *X*_**train**_): Reuse the temporal posteriors 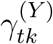 to re-estimate observation parameters 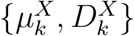 on *X*_train_ using their maximum-likelihood estimators.
3. **Step 3 (Temporal inference on** *X*_**test**_): With means and covariances 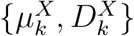 fixed, re-infer posterior time courses 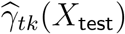. This is equivalent to retraining with `learn means=False` and `learn covariances=False` (see Supplementary Table S1), while updating all other parameters.
4. **Step 4 (Evaluation on** *Y*_**test**_): Combine 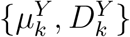 from step 1 with 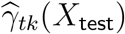 to compute the log-likelihood on *Y*_test_:

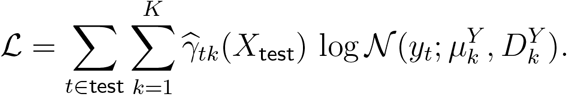

##### Notes

(1) When *K* = 1, steps 2 and 3 are omitted, and bi-cross-validation reduces to estimating a single covariance from *Y*_train_ and evaluating its log-likelihood on *Y*_test_. (2) In this paper, state means are not trained, so *µ*_*k*_ ≡ 0 throughout. (3) We tested whether binarising the posterior time courses *γ*_*tk*_ affected results, but found no substantial differences.

### A.2 Dynamic Network Modes (DyNeMo)

#### A.2.1 Generative model

Dynamic Network Modes (DyNeMo) (Gohil et al., 2022) extends the HMM by replacing mutually exclusive states with additive “modes.” At each time *t*, let *α*_*t*_ denote a vector of non-negative weights (summing to 1) over *M* modes, obtained by applying a softmax transformation to unconstrained latent variables *θ*_*t*_ ∈ ℝ^*M*^. The fMRI observation **x**_*t*_ is is assumed to follow

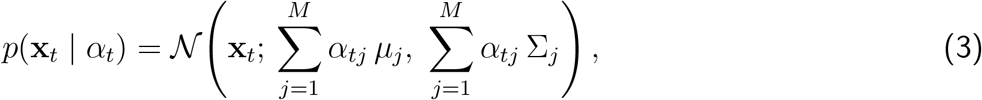

where each mode *j* is associated with a mean activation *µ*_*j*_ and covariance matrix Σ_*j*_.

The temporal dynamics of the latent coefficients *θ*_*t*_ are captured by a recurrent prior, implemented via a Long Short-Term Memory (LSTM) network:

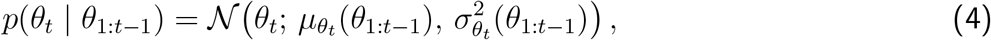

where 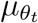 and 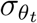 are neural networks mapping past coefficients to the parameters of a Gaussian distribution.

#### A.2.2 Inference

DyNeMo uses a variational Bayesian framework that minimises the variational free energy

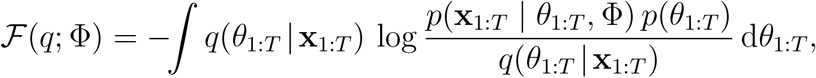

where 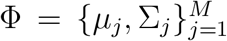 are the mode-specific observation parameters. Here, the likelihood *p*(**x**_1:*T*_ | *θ*_1:*T*_, Φ) depends only on Φ, while the prior *p*(*θ*_1:*T*_) captures the temporal structure of latent coefficients via an LSTM.

The approximate posterior is taken to be Gaussian at each time point,

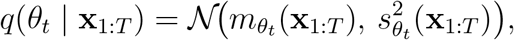

with mean 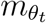 and variance 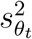 parameterised by a bidirectional LSTM.

Thus, DyNeMo inference yields a probabilistic description of the latent mode activations through *q*(*θ*_*t*_ | **x**_1:*T*_), while learning point estimates for the observation model parameters {*µ*_*j*_, Σ_*j*_}.

#### A.2.3 Bi-cross-validation procedure

Given a bi-cross-validation split, model evaluation proceeds as follows:

1. **Step 1 (Full train on** *Y*_**train**_): Estimate posterior mode activations *q*(*θ*_*t*_ | *Y*_train_) and observation parameters 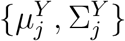 on *Y*_train_.
2. **Step 2 (Spatial inference on** *X*_**train**_): Reuse the temporal posteriors *q*(*θ*_*t*_ | *Y*_train_) to re-estimate the observation parameters 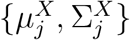 on *X*_train_. In practice, the posterior means 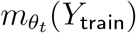 are used instead of full Gaussian samples when computing the maximum-likelihood estimates.
3. **Step 3 (Temporal inference on** *X*_**test**_): With observation parameters 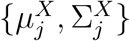 fixed, re-infer posterior mode activations 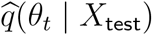 using the inference network. This is equivalent to retraining with `learn means=False` and `learn covariances=False` (see Supplementary Table S2), while updating all other parameters.
4. **Step 4 (Evaluation on** *Y*_**test**_): Combine 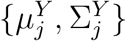 from Step 1 with the posterior means 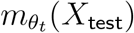 (rather than the full distributions 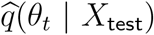 to compute the held-out log-likelihood on *Y*_test_.

##### Notes

(1) When *M* = 1, steps 2 and 3 are redundant, and bi-cross-validation reduces to estimating a single covariance from *Y*_train_ and evaluating its log-likelihood on *Y*_test_. (2) In this paper, mode means are not trained, so *µ*_*j*_ ≡ 0 throughout.

### A.3 Sliding Window Correlation (SWC)

#### A.3.1 Model procedure

Sliding window correlation (SWC) (Allen et al., 2014) characterises time-varying functional connectivity by computing covariances/correlations within short windows of the fMRI time series. Given fMRI data **x**_1:*T*_, we define windows of fixed length *L* with stride *δ* (window offset), producing *W* windows

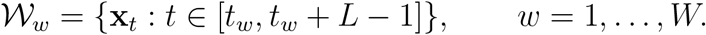

For each window *w*, we compute the sample covariance

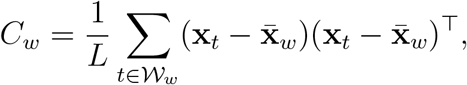

where 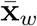 is the window mean. Each covariance *C*_*w*_ is then vectorised by extracting its upper-triangular entries, including the diagonal,

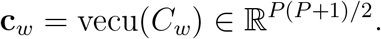

The vectors 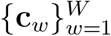 are clustered into *K* groups using *k*-means. Let *z*_*w*_ ∈ {1, …, *K*} denote the cluster label for window *w*, and let 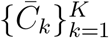 be the cluster centroids in matrix form:

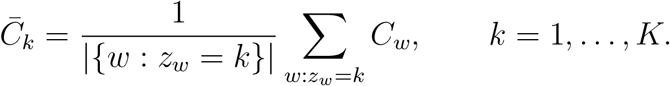

Since each centroid is the average of symmetric positive semidefinite matrices, it is itself symmetric positive semidefinite, and can therefore be interpreted as the covariance associated with state *k*.

Thus, the SWC procedure produces (i) a sequence of discrete labels {*z*_*w*_} describing the state assignment of each window, and (ii) state-specific covariance centroids 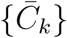 describing the spatial patterns of connectivity.

#### A.3.2 Bi-cross-validation procedure

Given a bi-cross-validation split, model evaluation proceeds as follows:

1. **Step 1 (Full train on** *Y*_**train**_): Compute window covariances {*C*_*w*_} on *Y*_train_, vectorise them to {**c**_*w*_}, and run *k*-means to obtain cluster centroids 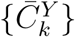 and labels 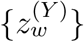.
2. **Step 2 (Spatial inference on** *X* _**train**_ ): Fix the cluster labels 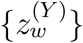 obtained on *Y* _train_, and recompute centroids 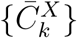 on *X*_train_ by averaging the corresponding window covariances.
3. **Step 3 (Temporal inference on** *X*_**test**_): Fix the centroids 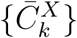 from Step 2, and reassign each window covariance from *X*_test_ to the nearest centroid, yielding new labels 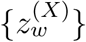.
4. **Step 4 (Evaluation on** *Y*_**test**_): Combine centroids 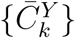 from Step 1 with window assignments 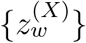 from Step 3 to compute the log-likelihood under the assigned centroid distributions on *Y*_test_.

##### Notes

(1) When *K* = 1, Steps 2 and 3 are redundant, and bi-cross-validation reduces to estimating a single covariance from *Y*_train_ and evaluating its fit to *Y*_test_. (2) Cluster labels are hard assignments (*z*_*w*_ ∈ {1, …, *K*}), in contrast to the probabilistic time courses of HMM and DyNeMo. (3) When the window length *L* spans the entire scanning session, the SWC procedure still applies, but each session contributes a single covariance matrix. In this case, the model reduces to clustering static functional connectivity across subjects.

**Table S1:**
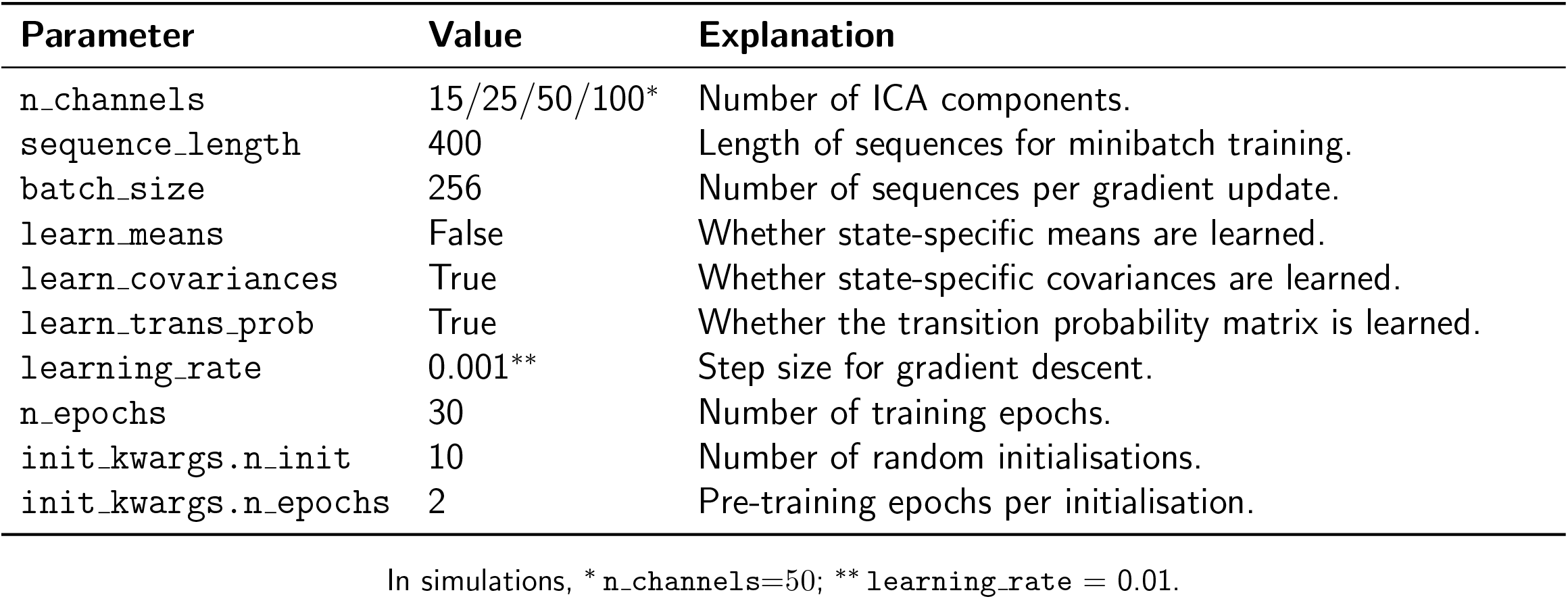
Hidden Markov Model (HMM) inference parameters.

**Table S2:** Dynamic Network Modes (DyNeMo) inference parameters.

| Parameter | Value | Explanation |
| --- | --- | --- |
| n_channels | 15/25/50/100* | Number of ICA components. |
| sequence_length | 100** | Length of sequences for minibatch training. |
| inference_n_units | 64 | Hidden units in inference network. |
| inference_normalization | layer | Normalisation in inference network. |
| model_n_units | 64 | Hidden units in dynamics network. |
| model_normalization | layer | Normalisation in dynamics network. |
| learn_alpha_temperature | True | Learn the softmax temperature controlling mode mixing. |
| initial_alpha_temperature | 1.0 | Initial value of the softmax temperature. |
| learn_means | False | Whether mode-specific means are learned. |
| learn_covariances | True | Whether mode-specific covariances are learned. |
| do_kl_annealing | True | Gradually increase KL penalty during training. |
| kl_annealing_curve | tanh | Functional form of KL annealing. |
| kl_annealing_sharpness | 5 | Steepness of annealing curve. |
| n_kl_annealing_epochs | 15 | Epochs for KL annealing. |
| batch_size | 64*** | Number of sequences per gradient update. |
| learning_rate | 0.001**** | Step size for gradient descent. |
| n_epochs | 30 | Number of training epochs. |
| init_kwargs.n_init | 10***** | Number of random initialisations. |
| init_kwargs.n_epochs | 2 | Pre-training epochs per initialisation. |

In simulations, \* n\_channels=50; \*\* sequence\_length = 200; \*\*\* batch\_size = 32; \*\*\*\* learning\_rate = 0.01;
\*\*\*\*\* init\_kwargs.n\_init = 50.

**Table S3:** Sliding window correlation (SWC) parameters.

| Parameter | Value | Explanation |
| --- | --- | --- |
| n_channels | 15/25/50/100* | Number of ICA components. |
| learn_means | False | Whether cluster/centroid means are learned. |
| learn_covariances | True | Whether cluster/centroid covariances are learned. |
| window_length | 100** | Sliding-window length (in TRs). |
| window_offset | 100 | Step size between successive windows (in TRs). |

Notes: \* n\_channels=50 in simulations; \*\* for the static FC model, window length equals the full session (1200 TRs for HCP; 490 TRs for UKB).

## B DyNeMo Additional Simulation

The main DyNeMo simulation in Section 2.4.2 used mode time courses and covariance matrices estimated from real HCP data. While this setup provides a realistic benchmark, the resulting simulated data inherit the unequal prominence of modes present in the trained model, with some modes being strong and persistent and others weak or short-lived. When evaluated with DyNeMo itself, this imbalance leads to conservative model behaviour: the bi-cross-validated log-likelihood increases up to four modes and then plateaus below the ground-truth value of six (Figure 3E), while the temporal recovery analysis reveals that only the more dominant modes are reliably reproduced (Figure 4).

To verify that this behaviour reflects the intrinsic data structure and training characteristics of DyNeMo rather than a limitation of bi-cross-validation, we conducted a supplementary DyNeMo control simulation using the same basic configuration: six modes shared across 500 subjects, each with 1200 time points. The key difference lay in the data generation procedure. Instead of using modes derived from real data, we generated both the mode time courses and mode covariance matrices synthetically. Mode time courses were defined as sinusoidal functions with varying frequencies and phases, whereas mode covariances were sampled as random symmetric positive-definite (SPD) matrices following the approach described in the original DyNeMo paper (Gohil et al., 2022).

More specifically, for each mode *k*, an unnormalised activation trace was defined as

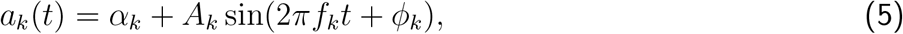

where *α*_*k*_ controls the baseline level, *A*_*k*_ the amplitude, *f*_*k*_ the frequency, and *ϕ*_*k*_ ~ 𝒰(0, 2*π*) a random phase offset. These activations were then normalised across modes at each time point using a softmax function,

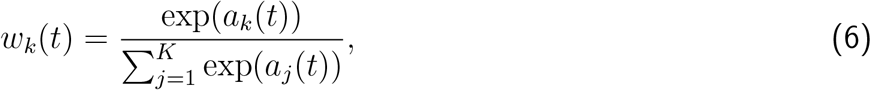

so that the weights *w*_*k*_(*t*) represent smoothly varying mode activations that sum to one.

Covariance matrices Σ_*k*_ were independently drawn as random symmetric positive-definite (SPD) matrices to represent distinct spatial patterns of co-fluctuation. Specifically, random weight matrices *W*_*k*_ ∈ ℝ^*R*×*R*^ were sampled from a normal distribution *W*_*k*_ ~ 𝒩 (0, 0.1^2^), and covariances were computed as

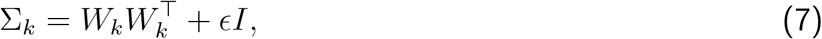

where a small diagonal term *ϵI* (with *ϵ* = 10^−6^) ensured numerical stability. To introduce mild structure, a random subset of channels received slightly higher weights during the construction of *W*_*k*_, producing non-isotropic but well-conditioned covariance patterns. The full set of simulation parameters used in this control analysis is listed in Table S4.

Using this fully synthetic DyNeMo dataset, we then applied the same bi-cross-validation procedure as in the main simulation to evaluate HMM, DyNeMo, and SWC across a range of model orders. Figure S2 summarises the resulting bi-cross-validated log-likelihood profiles.

For DyNeMo, the bi-cross-validated log-likelihood increases up to seven modes (log-likelihood = 1.149 ± 0.157). The dagger at seven modes identifies a model order close to the ground-truth value of six. In contrast, HMM performance continued to increase across the tested range, peaking at sixteen states (log-likelihood = 1.084 ± 0.042), but remained suboptimal relative to DyNeMo. SWC again showed the weakest performance overall, with substantially lower log-likelihood values and a peak at thirteen states (log-likelihood=0.092 ± 0.011). Together, these results confirm that the conservative behaviour in the main DyNeMo simulation reflects properties of the data and model interaction rather than a limitation of bi-cross-validation itself.

**Figure S1:**
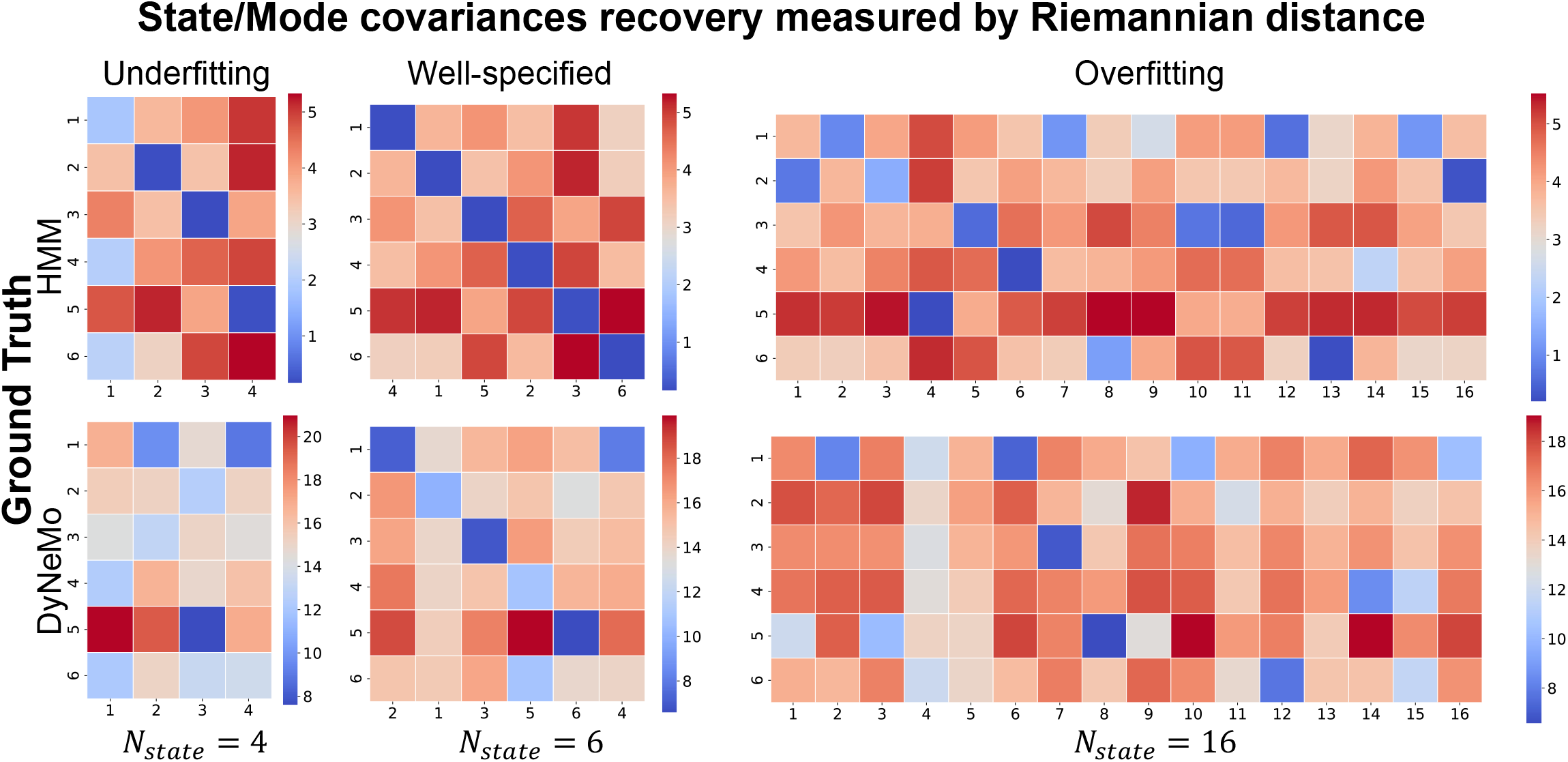
For the HMM 6-state simulation (first row), we trained three HMMs with 4 states (under-fitting), 6 states (well-specified), and 16 states (overfitting) on the full dataset. We then computed the Riemannian distance between the inferred and ground-truth state covariances. For the well-specified model (6 states), state correspondence was optimised using the Hungarian algorithm. The second row shows the same procedure applied to the 6-mode DyNeMo simulation.

**Table S4:** Parameters used in the DyNeMo additional control simulation implemented via simulation.MixedSine MVN in the osl-dynamics toolbox.

| Parameter | Value | Description |
| --- | --- | --- |
| n_modes | 6 | Number of simulated modes. |
| n_channels | 50 | Number of spatial channels (brain regions). |
| n_samples | $500 \times 1200$ | Total number of time points across all subjects. |
| relative_activation | [1.0, 1.75, 2.5, 3.25, 4.0, 4.75] | Baseline activation levels $\alpha_k$ in Eq. 5. |
| amplitudes | [6, 5, 4, 3, 2, 1] | Amplitudes $A_k$ controlling oscillation strength in Eq. 5. |
| frequencies | [1.3, 2.3, 3.3, 4.3, 5.3, 6.3] Hz | Mode oscillation frequencies $f_k$ in Eq. 5. |
| sampling_frequency | 250 Hz | Sampling rate for sinusoidal time-course generation. |
| means | “zero” | Mean vector for each mode set to zero. |
| covariances | “random” | Covariances generated as random SPD matrices via Eq. 7. |
| n_covariances_act | 3 | Number of activation iterations for covariance construction, introducing structured variability. |
| epsilon | $10^{-6}$ | Diagonal regularization term ensuring positive definiteness of $\Sigma_k$. |

**Figure S2:**
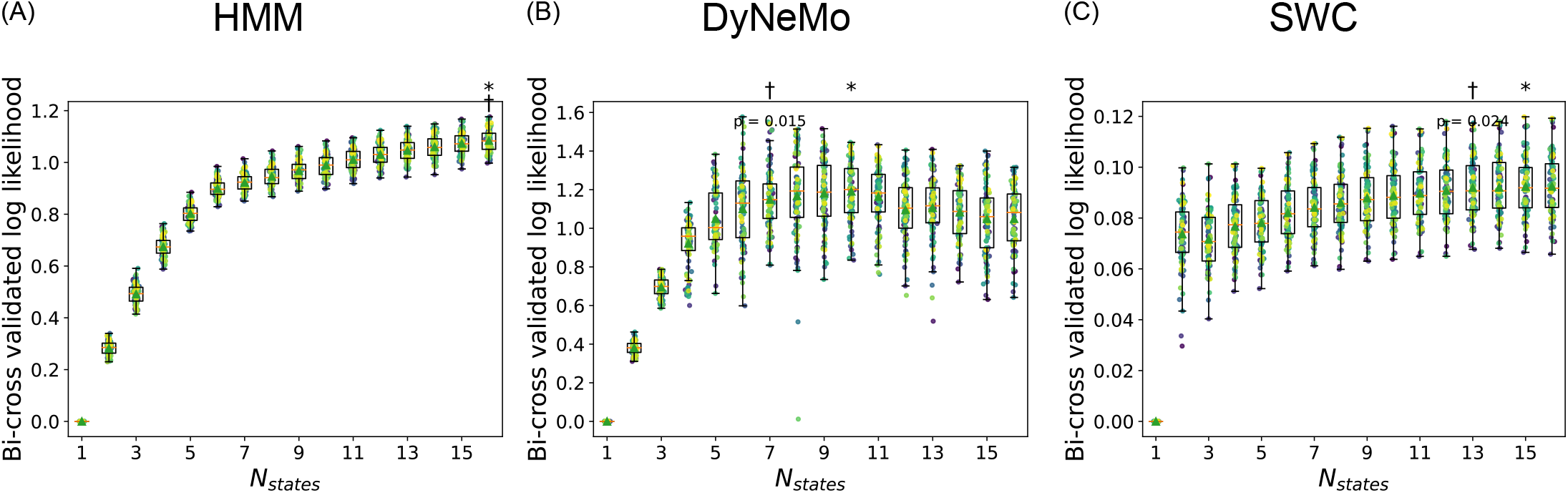
Dynemo Additional Simulation Results. Bi-cross-validation results for the supplementary DyNeMo control simulation with a ground-truth of six modes. Panels show evaluations of (A) HMM, (B) DyNeMo, and (C) SWC. In each panel, the x-axis indicates the number of states or modes used during model fitting, and the y-axis shows the bi-cross-validated log-likelihood. Each dot represents a single bi-cross-validation realisation, demeaned by the corresponding static model baseline (i.e., *N*_states_ = 1). Dots of the same color across values of *N*_states_ correspond to the same “reshuffle-and-split” realisation. Box plots show the distribution of log-likelihoods across realisations. An asterisk indicates the best-performing hyperparameter in each panel. A dagger marks the smallest number of states or modes whose performance is not significantly worse than the best-performing model, based on a one-sided paired t-test (*p <* 0.001).

**Figure S3:**
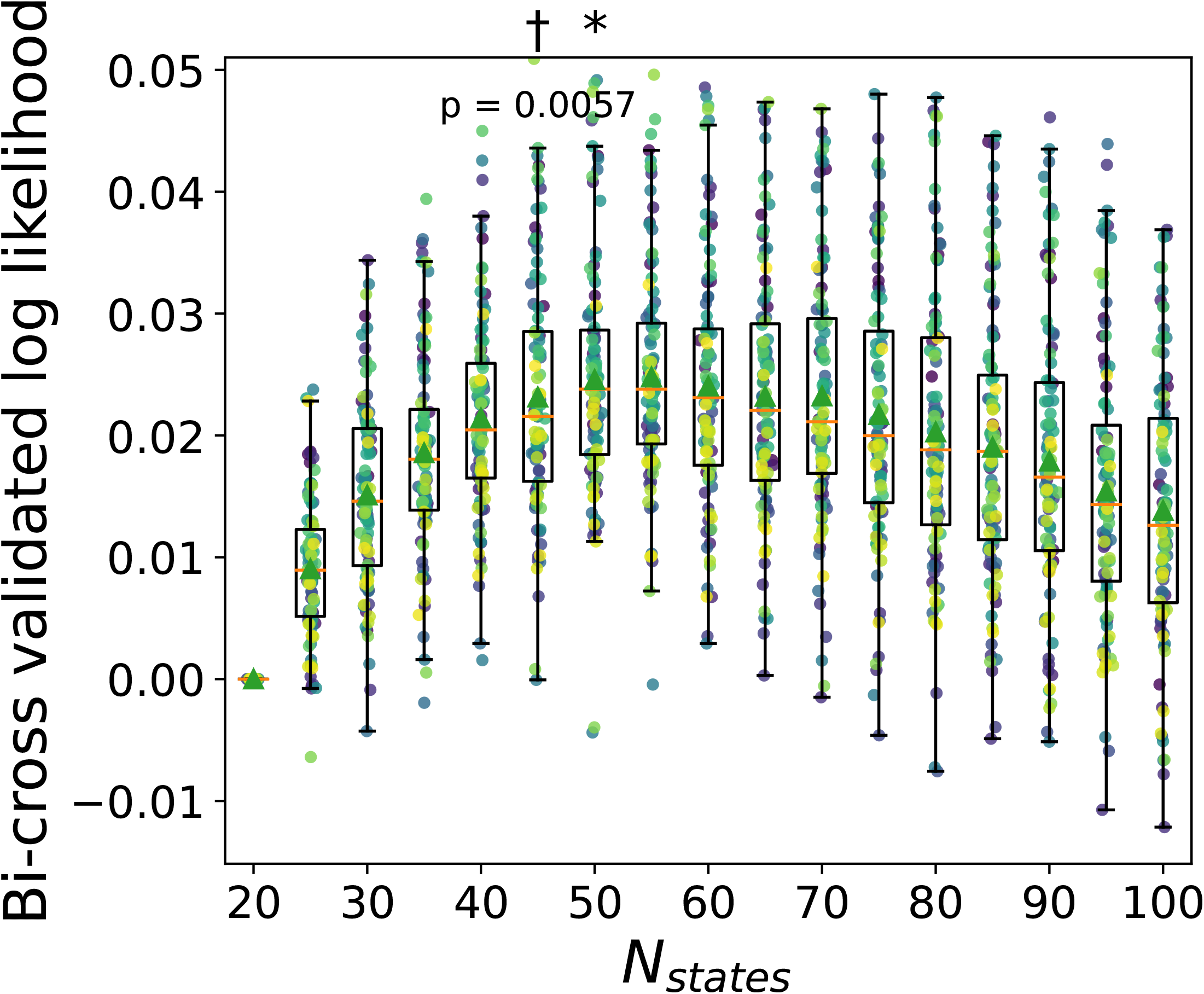
SWC’s performance on HCP ICA 50. Resting-state fMRI data from the HCP were processed using group-level spatial ICA with 50 components, followed by dual regression to extract subject-level time series. Bi-cross-validation was applied to SWC across model orders ranging from 20 to 100. All plots were generated using the same procedure as in Figure 5 except that log-likelihoods were demeaned relative to the SWC model with 20 states. The asterisks mark the model with the highest median log-likelihood, and daggers indicate the smallest model not significantly worse than the best (one-sided paired t-test, *p <* 0.001).

**Figure S4:**
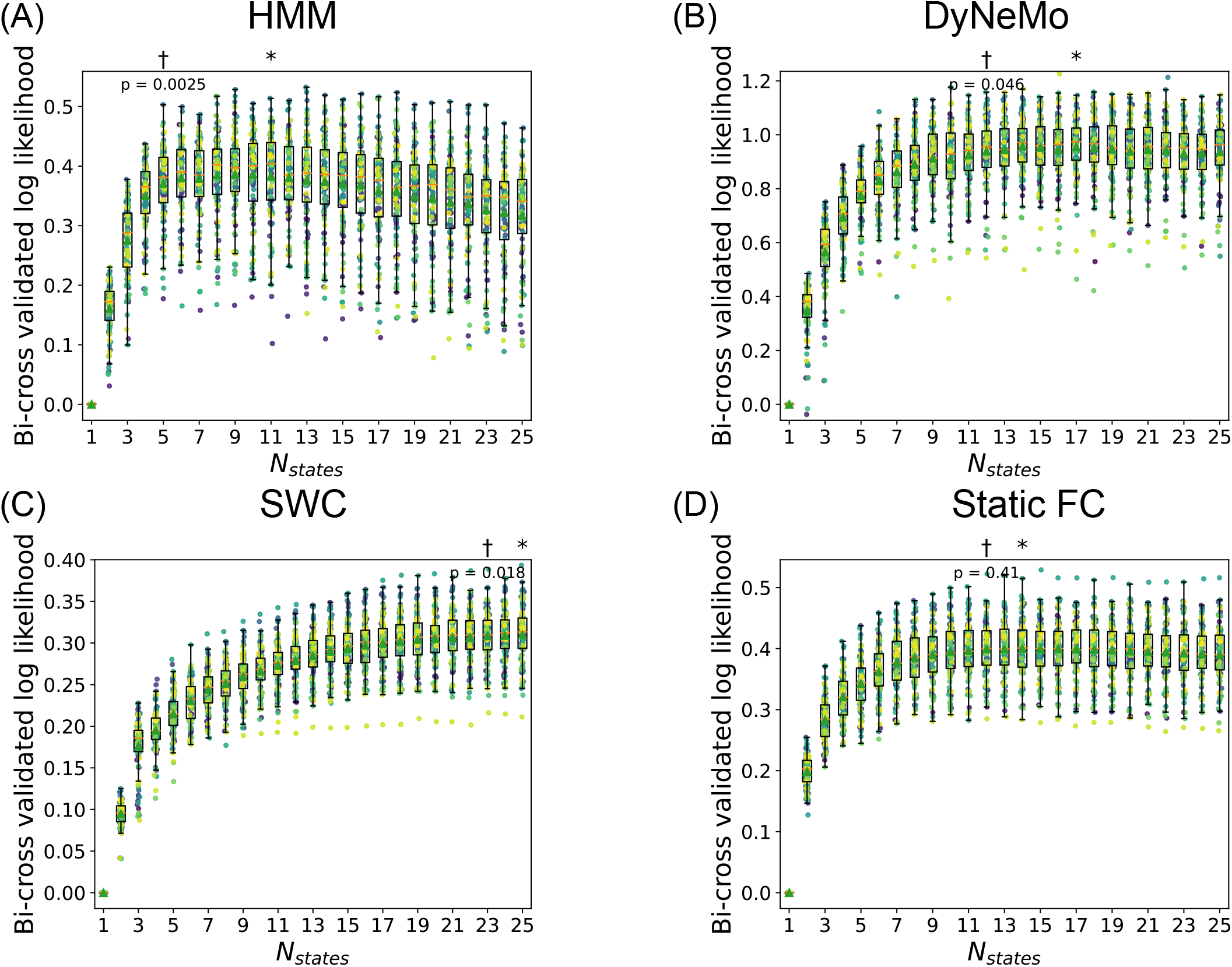
Model performance on UKB ICA 50. (A) HMM, (B) DyNeMo, (C) SWC (D) static FC. Resting-state fMRI data from the UKB dataset were processed using group-level spatial ICA with 50 components, followed by dual regression to obtain subject-level time series. Bi-cross-validation was applied to each model with number of states from 1 to 25. All plots were generated using the same procedure as in Figure 5, including asterisks to indicate the highest median log-likelihood and daggers to denote the smallest model not significantly worse than the best (one-sided paired t-test, *p <* 0.001).

**Figure S5:**
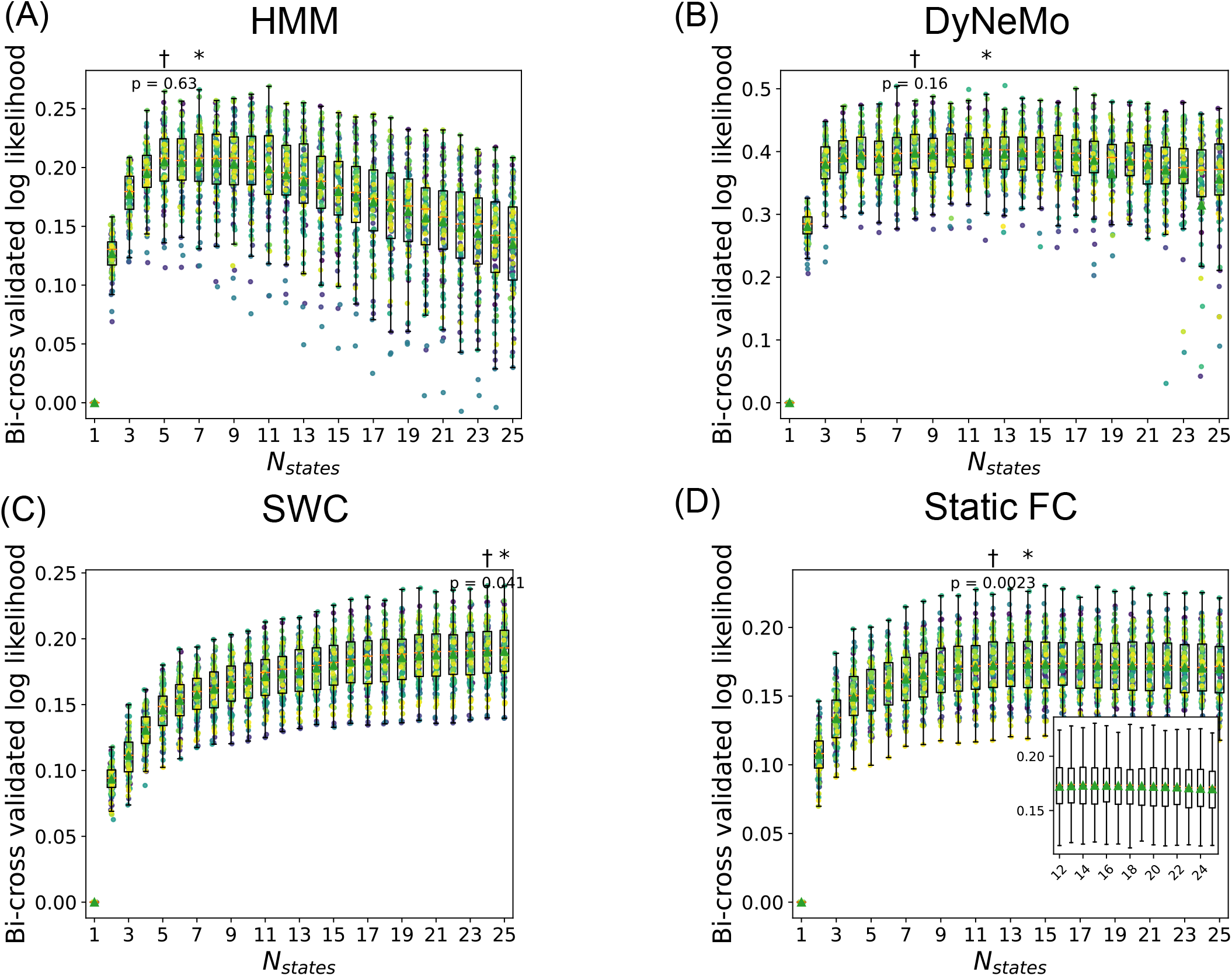
Model performance on HCP ICA 50 high-pass filtered data. (A) HMM, (B) DyNeMo, (C) SWC (D) static FC. Resting-state fMRI data from the HCP were processed using group-level spatial ICA with 50 components, followed by dual regression to extract subject-level time series. The resulting time-series were high-pass filtered using a fifth-order Butterworth filter with a cut-off frequency at 0.05 Hz. Bi-cross-validation was applied to each model across model orders ranging from 1 to 25. All plots were generated using the same procedure as in Figure 5, with asterisks marking the model with the highest median log-likelihood, and daggers indicating the smallest model not significantly worse than the best (one-sided paired t-test, *p <* 0.001). The inset in panel (D) provides a zoomed-in view over *N*_states_ = 12 to 25 to highlight differences in model performance.

**Figure S6:**
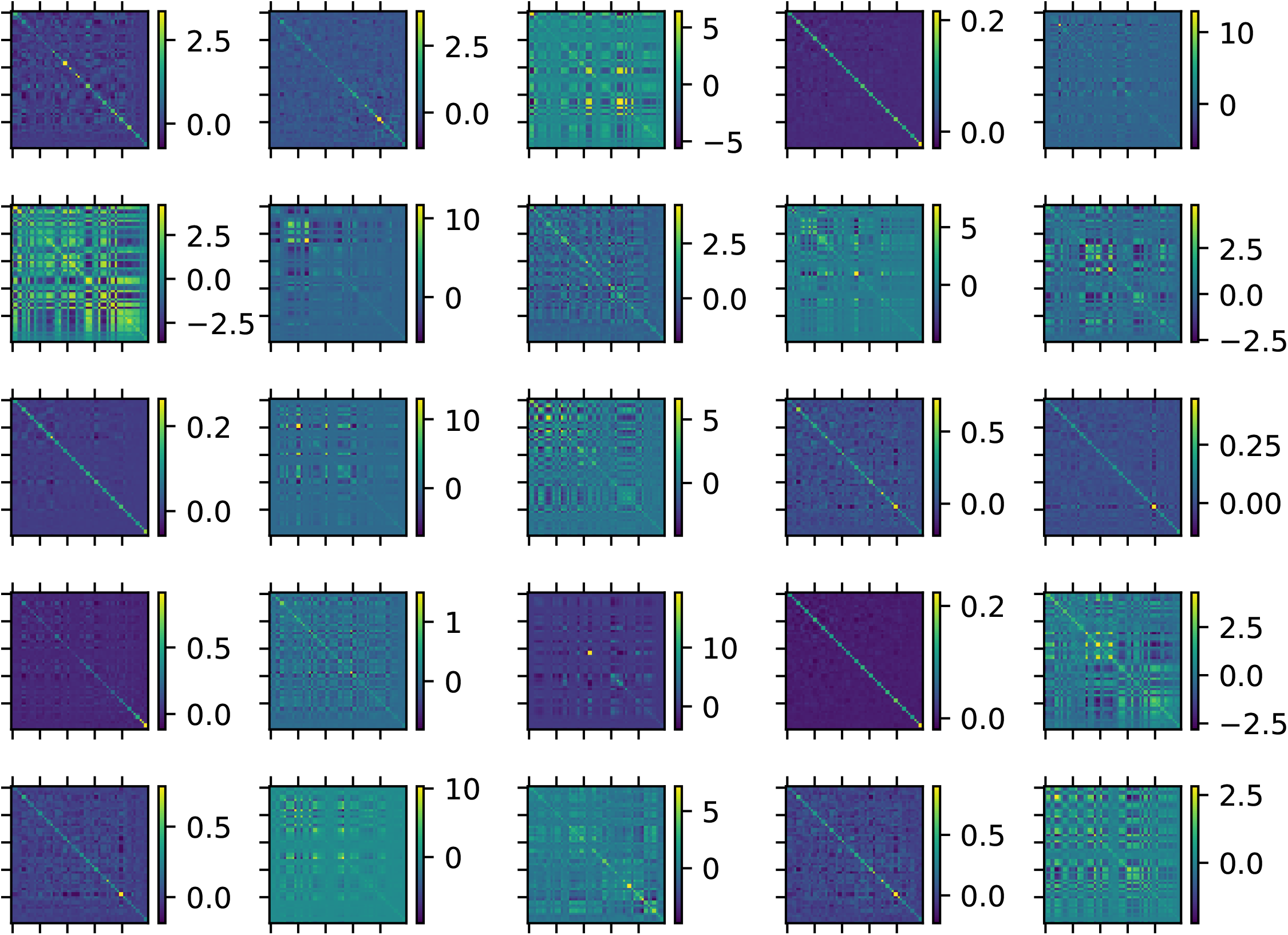
Mode covariances of 25-mode DyNeMo on HCP ICA 50. Resting-state fMRI data from the HCP were processed using group-level spatial ICA with 50 components, followed by dual regression to extract subject-level time series. A DyNeMo model with 25 modes (zero-mean activation) was trained on the full dataset, and the resulting mode covariances are visualised. The least-activated modes (e.g., modes 4,11,19,15) closely resemble diagonal matrices due to the prior distribution imposed on mode covariances.

**Figure S7:**
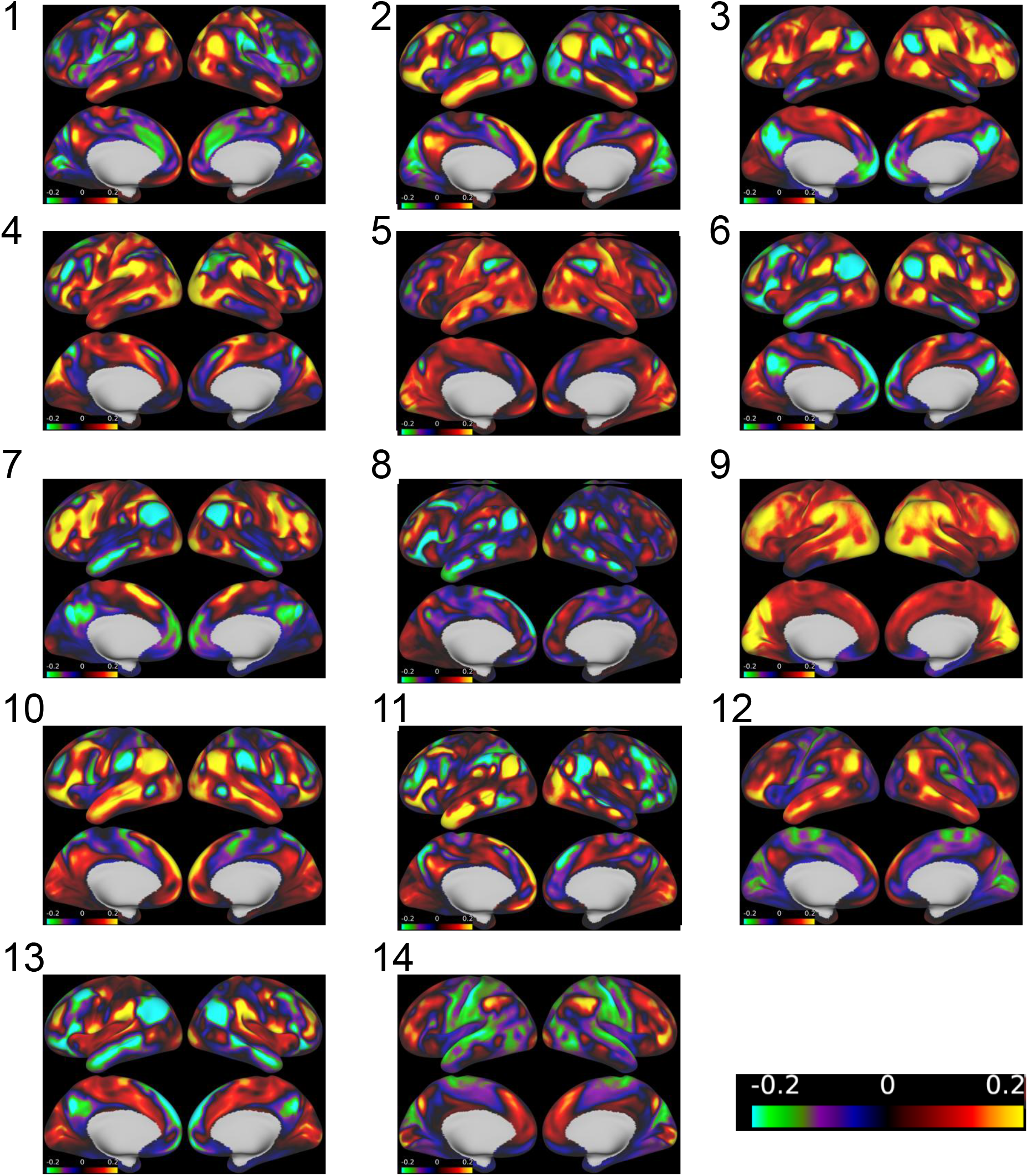
Visualisation of full DyNeMo modes (HCP ICA 50). A 14-mode DyNeMo model was trained on HCP ICA 50. After training, mode covariance matrices were converted to correlation matrices. For each mode, a rank-one approximation was computed by extracting the dominant eigenvector (i.e., the eigenvector associated with the largest eigenvalue) of its mode correlation matrix. The resulting eigenvector was then projected back to brain space to obtain a spatial map for visualisation. A single symmetric colour scale centred at zero was used for all states and modes.

**Figure S8:**
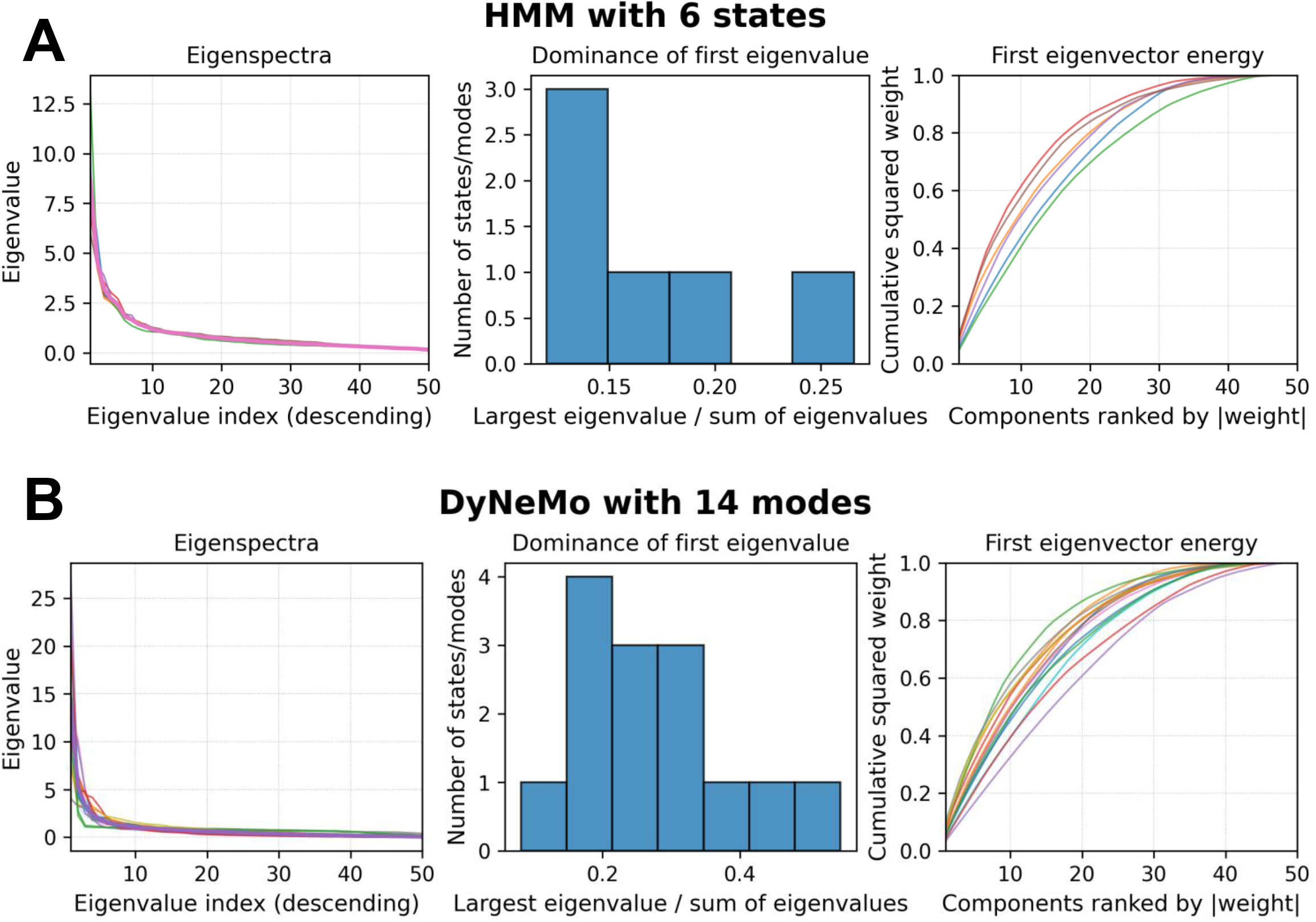
Eigenspectrum of HMM states and DyNeMo modes (HCP ICA 50). A 6-state HMM (A) and a 14-mode DyNeMo model (B) were trained on HCP ICA 50. After training, state or mode covariance matrices were converted to correlation matrices. For each state or mode, the eigenvalues of the corresponding correlation matrix are shown in descending order (left), where each line represents one state (HMM) or mode (DyNeMo). The histogram (middle) shows, for each state or mode, the ratio between the largest eigenvalue and the sum of all eigenvalues. The cumulative curves (right) show the cumulative sum of squared weights of the first eigenvector after sorting ICA components by the absolute value of their weights.

**Figure S9:**
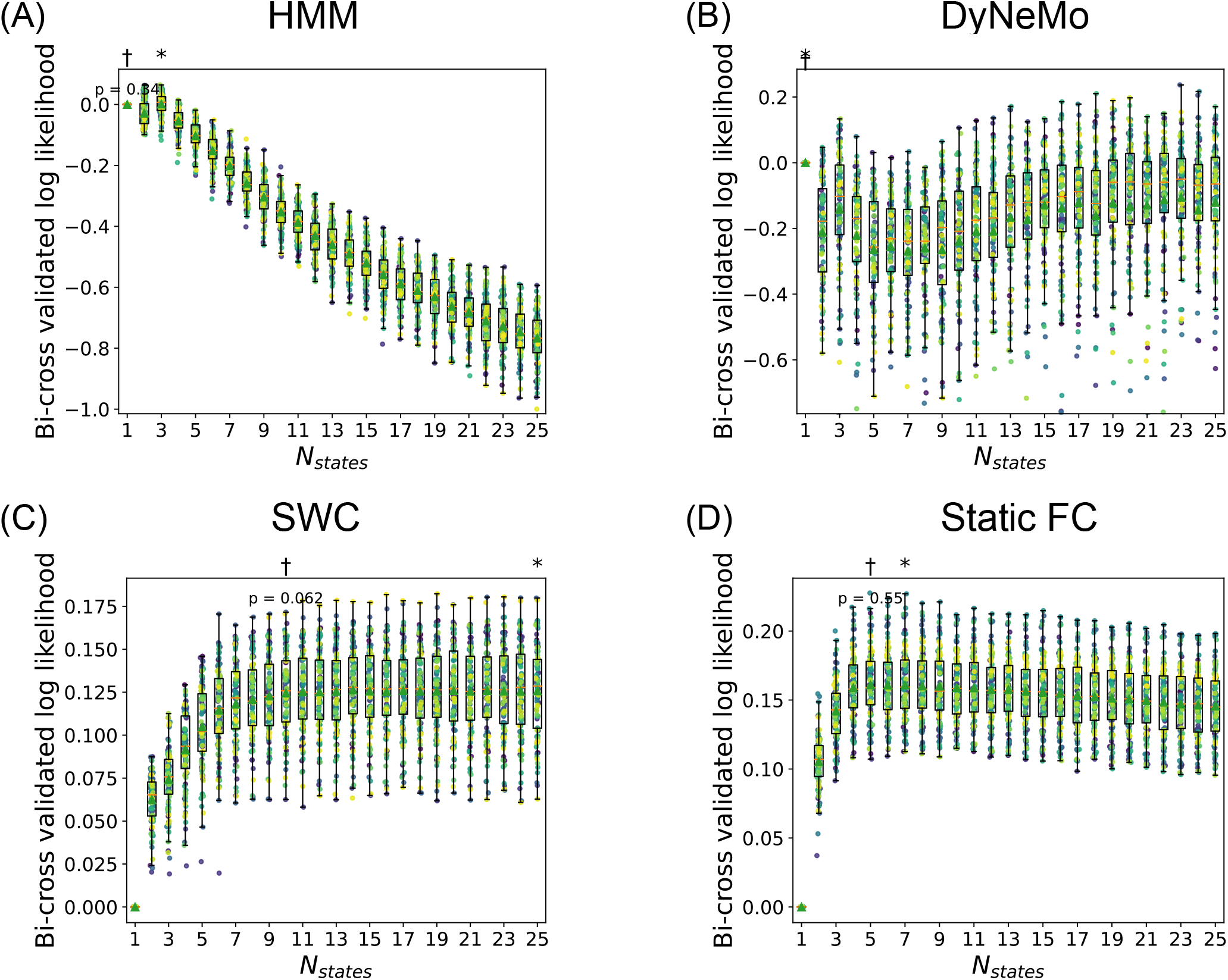
Model performance on HCP ICA 25. (A) HMM, (B) DyNeMo, (C) SWC (D) static FC. Resting-state fMRI data from the HCP were processed using group-level spatial ICA with 25 components, followed by dual regression to extract subject-level time series. Bi-cross-validation was applied to each model across model orders ranging from 1 to 25. All plots were generated using the same procedure as in Figure 3, with asterisks marking the model with the highest median log-likelihood, and daggers indicating the smallest model not significantly worse than the best (one-sided paired t-test, *p <* 0.001).

**Figure S10:**
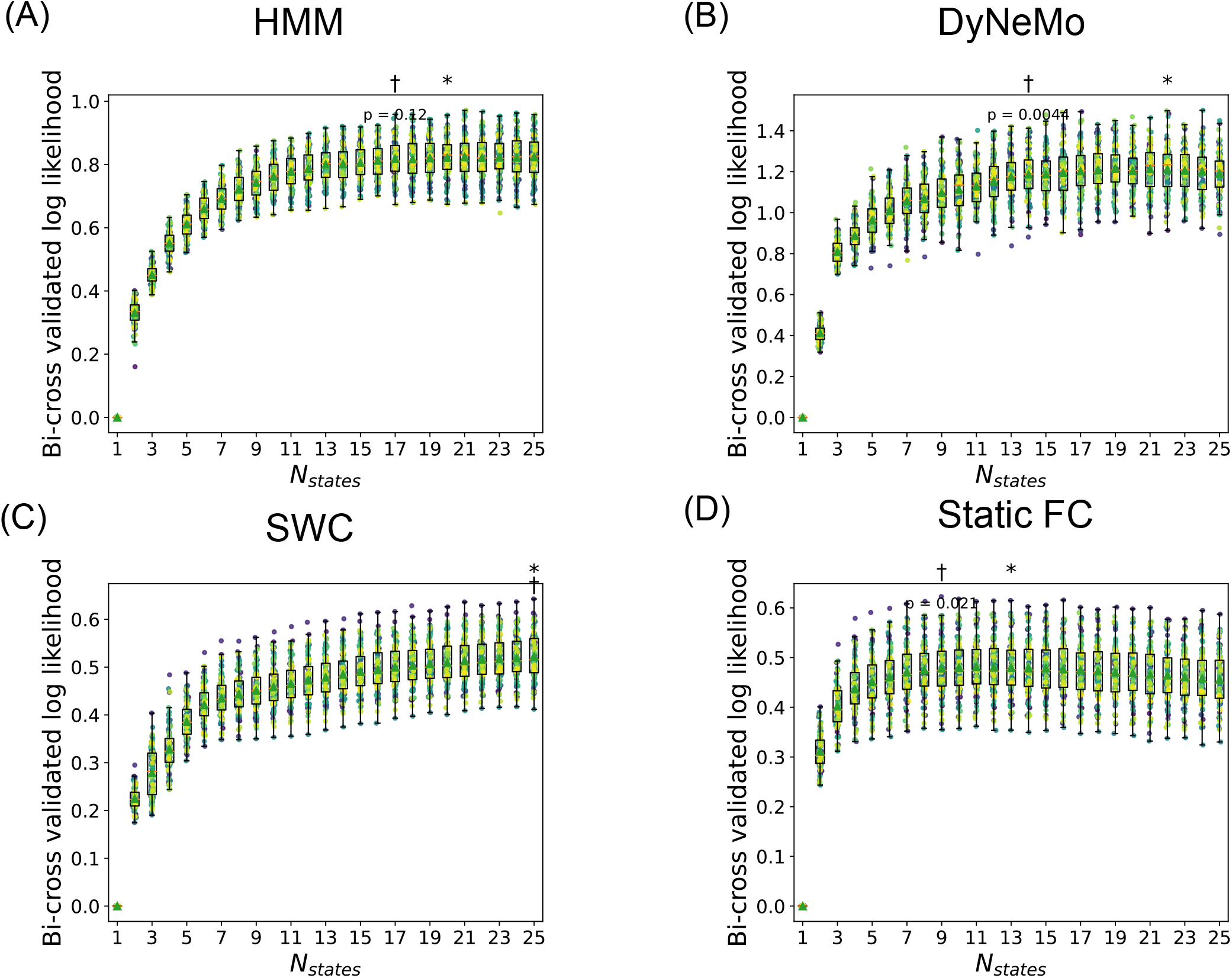
Model performance on HCP ICA 100. (A) HMM, (B) DyNeMo, (C) SWC (D) static FC. Resting-state fMRI data from the HCP were processed using group-level spatial ICA with 100 components, followed by dual regression to extract subject-level time series. Bi-cross-validation was applied to each model across model orders ranging from 1 to 25. All plots were generated using the same procedure as in Figure 3, with asterisks marking the model with the highest median log-likelihood, and daggers indicating the smallest model not significantly worse than the best (one-sided paired t-test, *p <* 0.001).

**Figure S11:**
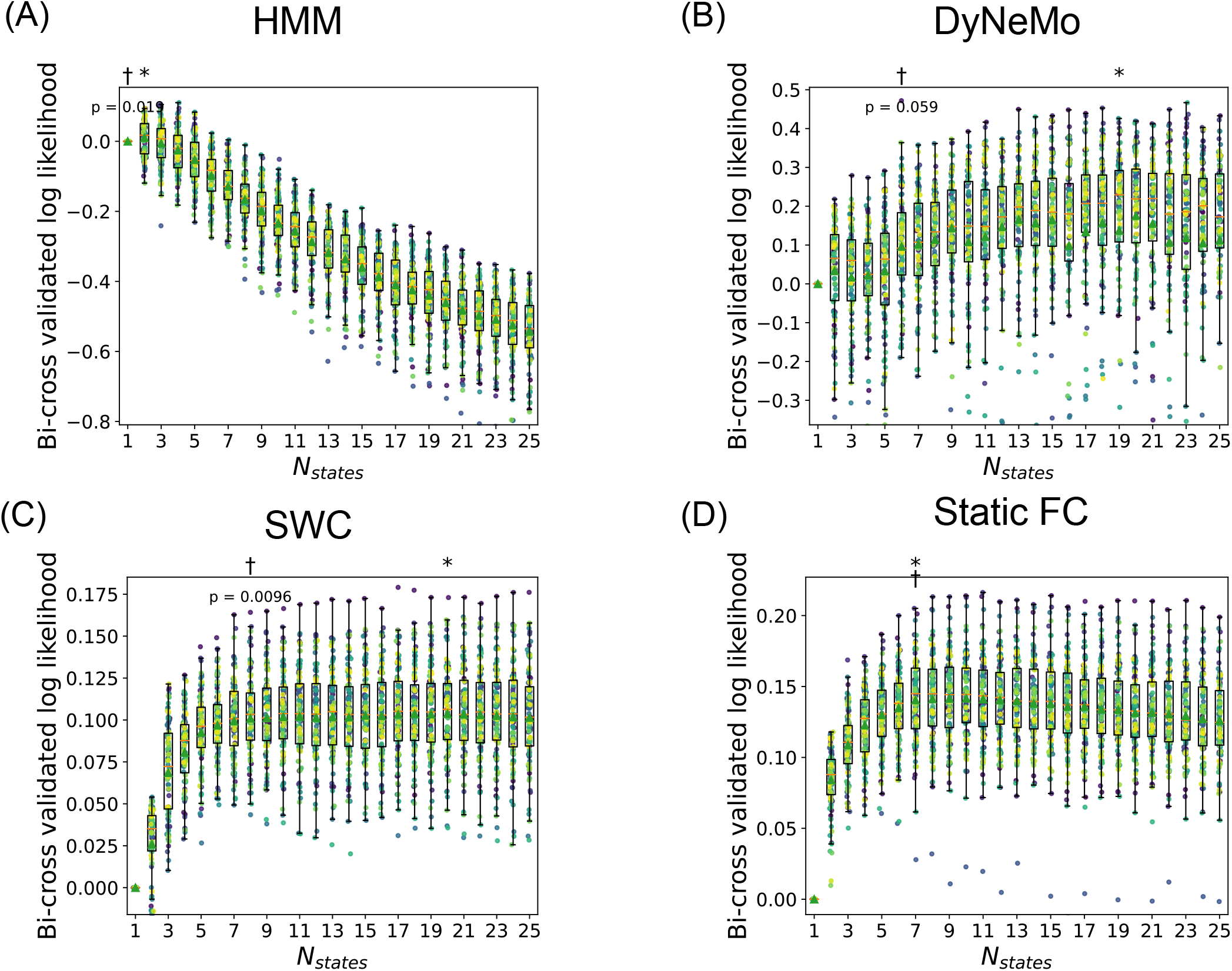
Model performance on UKB ICA 25. (A) HMM, (B) DyNeMo, (C) SWC (D) static FC. Resting-state fMRI data from UKB dataset were processed using group-level spatial ICA with 25 components, followed by dual regression to obtain subject-level time series. Bi-cross-validation was applied to each model with number of states from 1 to 25. All plots were generated using the same procedure as in Figure 5, including asterisks to indicate the highest median log-likelihood and daggers to denote the smallest model not significantly worse than the best (one-sided paired t-test, *p <* 0.001).

## Footnotes

1 Strictly speaking, the states in DyNeMo and MAGE are better described as “modes”, reflecting their additive combination in the generative model. However, for simplicity, they are sometimes referred to as states in this paper.

